# The tumor suppressor role of the *KRAS* proto-oncogene in metastatic pancreatic cancer

**DOI:** 10.64898/2026.11.18.767567

**Authors:** Li Zhang, Enrico Gurreri, Haoyu Tang, Angelina Bania, Mohammad J. Ibrahim, Roberta Rinaldi, Aastha Pandey, Luca Cecchetto, Elisabetta Granato, Hania Khan, Anh Hoang, Desiree Catania, Calogero Carlino, Joseph R Marszalek, Chunhua Shi, Alessandro Sgambato, Giampaolo Tortora, Alessandro Carugo, Andrea Viale, Davide Cittaro, Stefano Sioletic, Laura Wood, David S Klimstra, Olca Basturk, Shubham Pant, Milind Javle, Pavlos Msaouel, Jianjun Gao, Kadir C Akdemir, Linghua Wang, Peter Van Loo, Francesca Citron, Timothy Heffernan, Christine A. Iacobuzio-Donahue, Scott Kopetz, Luigi Perelli, Giannicola Genovese

## Abstract

*KRAS* is the most frequently mutated proto-oncogene in human cancers^1–3^. Progression to advanced disease is frequently associated with allelic imbalance of the *KRAS* locus with gain of the mutant allele and loss of the wild-type allele through clonal and sub-clonal loss of heterozygosity (LOH)^2^. While a large body of literature has focused on the effects of allelic imbalance on oncogene addiction through gain of the mutant allele, the role of the wild-type *KRAS* proto-oncogene in malignant progression has not been fully elucidated.

To capture this phenomenon, we developed a first-in-class somatic mosaic and lineage tracing genetically engineered mouse model (smGMLT) of pancreatic ductal adenocarcinoma allowing for real time *in vivo* tracing of *Kras*-LOH. Longitudinal characterization of the smGMLT demonstrated the presence of a strong evolutionary bottleneck towards the selective loss of the wild-type allele at metastatic sites. Cross species analysis leveraging research autopsies and surgical specimens revealed that *KRAS*-LOH is associated with metastatic dissemination and shorter overall survival. Transcriptomic analysis through single-cell RNA sequencing uncovered a conserved phenomenon whereby *KRAS*-LOH is paralleled by an immune evasive phenotype through loss of the tumor-cell interferon response and reduced CD8+ T cell infiltration.

In line with this evidence, orthotopic transplants of isogenic tumor populations differing for the *Kras* allelic status demonstrated that only cancer cells with *Kras*-LOH are are able to form tumors in immune competent animals. Restoration of the wild-type allele abrogates tumorigenesis in *Kras*-LOH transplants in immunocompetent through activation of the TBK1-IRF axis.

Altogether, these studies demonstrate that the *KRAS* proto-oncogene is a functional tumor suppressor in pancreatic cancer exerting its function through interferon-mediated, cell-extrinsic modulation of the tumor immune microenvironment.

## Introduction

Activating driver mutations of the *KRAS* oncogene account for about 20% of all human malignancies and more than 90% of pancreatic cancer^1–3^. Heterozygous single nucleotide variations (SNVs) resulting in impaired GTP hydrolysis and enhanced protein activity are sufficient to promote unrestrained cellular proliferation and enable malignant transformation^4–6^. However, accumulating genomic evidence indicates that the magnitude of oncogenic signaling is not uniquely determined by the mutational status, but further modulated by the relative oncogenic dosage and stoichiometry of mutant and wild-type alleles^2,5,7–9^. These phenomena are highly dynamic as several evolutionary bottlenecks converge on the *KRAS* locus. Genetic mechanisms altering the balance of mutant and wild-type alleles, including loss-of-heterozygosity (LOH), copy-number gain of the mutant allele, whole-genome doubling (WGD), and allele-specific transcriptional or epigenetic regulation, have been identified across tumor types through next generation sequencing (NGS) studies and are increasingly implicated as recurrent, selected events during clonal evolution rather than passenger alterations^2,5,8–10^. *KRAS*-mutant pancreatic cancers provide a remarkable example of this principle with immediate clinical implications. Clinical and genomic analysis of large patient cohorts has shown that changes of *KRAS* allelic copy number configurations are hallmarks of *KRAS*-mutant pancreatic ductal adenocarcinoma (PDAC) and correlate with clinical and pathological parameters of disease aggressiveness^2^.

These findings support a model in which evolutionary pressures on *KRAS* dosage, shape clinical trajectories, and potentially, response to therapy. Despite this progress, the spatial and temporal dynamics of allele selection at the *KRAS* locus during tumor evolution, and the functional consequences for cancer cells, remain incompletely defined. It is unclear whether allelic selection at the *KRAS* locus represents an early driver event in tumorigenesis or a later adaptative phenomenon that emerges only at advanced stages when additional selective pressures are imposed^5,8,11^.

From a functional standpoint, allelic imbalance at the *KRAS* locus is a result of the intrinsic oncogene addiction of cancer cells initiated by such a driver event^5,8^. Oncogene-addicted cells are, by definition, reliant on sustained pathway output for their proliferative and survival phenotype, to the extent that acute inhibition or genetic loss of the driver oncogene is sufficient to trigger growth arrest or apoptosis even in the presence of a fully transformed, genomically complex background^12,13^. This dependency implies that the tone of oncogenic signaling of a cancer cell is itself under continuous selective pressure, therefore supporting the hypothesis that oncogene allelic imbalance serves as an accelerator to gain a further fitness advantage from any mechanism that increases oncogenic tone. Under this model, allelic selection is not a passive consequence of genomic instability, but an active evolutionary strategy through which tumor cells titrate oncogenic signaling upward to maintain, or further reinforce, their addiction to the driver pathway.

To gain functional insight into the timing and functional consequences of allele selection in *KRAS*-mutant cancers, we generated a set of tools and cross-species analysis enabling us to trace and perform functional studies on *KRAS* allelic status during PDAC evolution leveraging in vivo somatic mosaic genome editing and lineage tracing (smGMLT).

### *KRAS* LOH is selected during metastatic progression in PDAC

Oncogene dynamics and allelic selection at the *KRAS* locus were first characterized through cross-species clinicogenomic analysis of PDAC leveraging surgical (N = 2,336) and autopsy (N = 30) samples from patient cohorts and pre-clinical models (N = 590) (^2,5,142,5,14^. This analysis shows that *KRAS*-LOH isa ssociated with aggressive disease, whole genome doubling, and metastatic dissemination across cohorts, independently of the *KRAS* driver mutation (**Figure 1 a-c, Extended data Figure 1 a-f**). End-stage disease, as studied through a series of research autopsies, demonstrates a significant enrichment of *KRAS* LOH at metastatic sites samples, suggesting that loss of the wild-type allele is positively selected in disseminated disease and during the emergence of clones with metastatic competencies (**Figure 1 d-e**).

**Figure 1:**
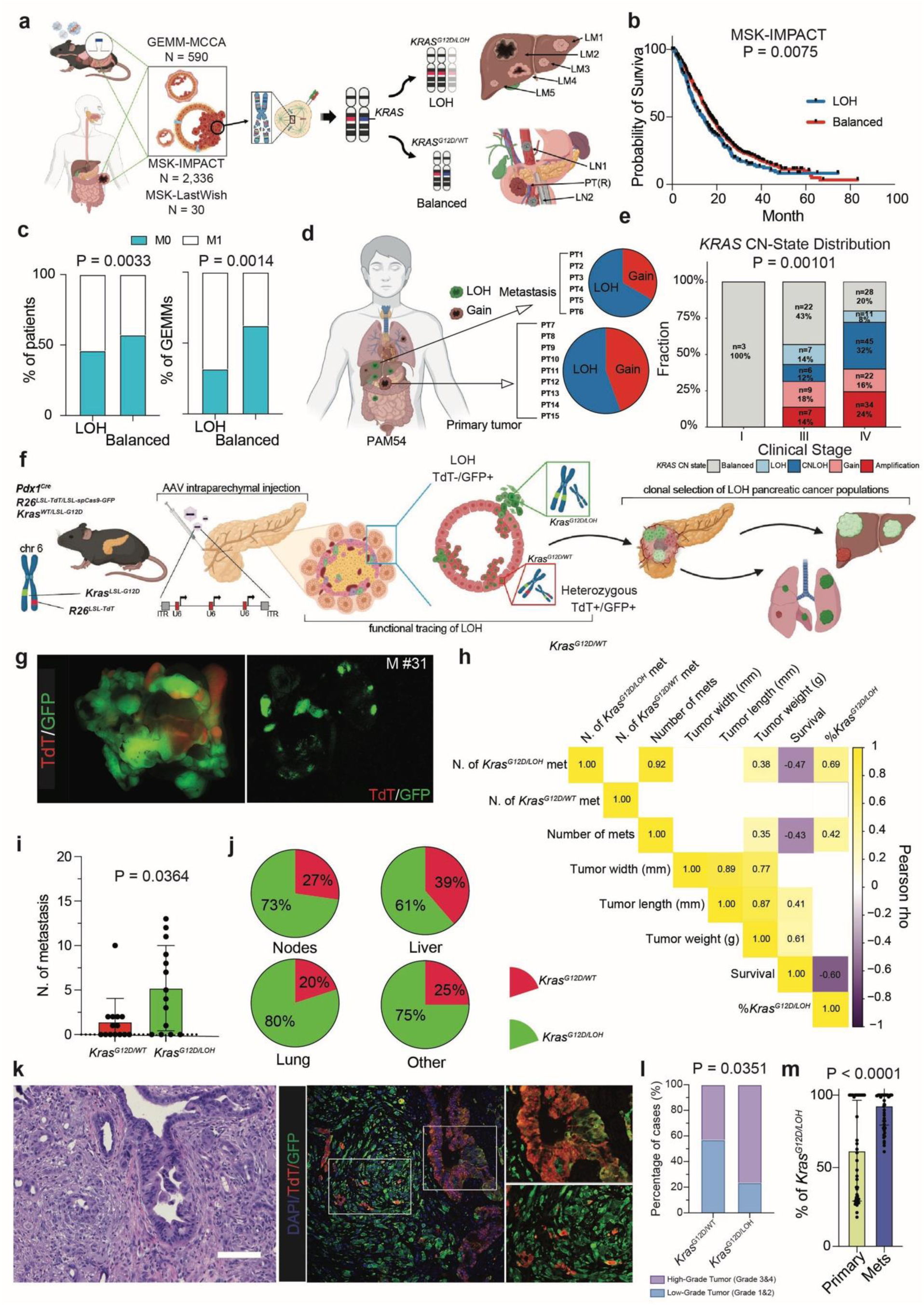
*KRAS* LOH is selected during metastatic progression in PDAC. **a**, Study design integrating GEMM model data (GEMM-MCCA)^5^ with patient cohorts (MSK-IMPACT, MSK-LastWish)^2,28^ to trace malignant lineages with mutant *KRAS* LOH or a balanced allelic state across primary and metastatic sites. **b**, Overall survival of MSK-IMPACT patients (N = 923) stratified by *KRAS* LOH (N = 242) versus balanced (N = 681) allelic state (*P* = 0.0075). **c**, Proportion of non-metastatic (M0, N = 791) and metastatic (M1, N = 1,416) cases among patients (left) and (M0, N = 37; M1, N = 55) SM-GEMMs (right) by allelic status (*P* = 0.0033 and *P* = 0.0014, respectively). **d,** Representative case of rapid-autopsy (PAM54) analysis of *KRAS* allelic status across matched metastatic (PT1–PT6) and primary (PT7–PT15) lesions. **e**, Distribution of *KRAS* copy-number states of MSK rapid-autopsies cohort (Balanced, LOH, copy-neutral LOH, Gain, Amplification) across clinical stage (I, III, IV); *n* per group as indicated (*P* = 0.00101)^28^. **f**, Schematic of the *K^LOH^C* SM-GEMM: the *Pdx1-Cre*, *R26^LSL-TdT/LSL-spCas9-GFP^* and *Kras^G12D/WT^* alleles, the configuration of the *Kras* and *Rosa26* loci on chromosome 6, AAV intraparenchymal delivery, and functional tracing of LOH, resolving *Kras^G12D/LOH^* (TdT^−^/GFP^+^) and heterozygous *Kras^G12D/WT^* (TdT^+^/GFP^+^) populations and their clonal selection. **g**, Representative macroscopic fluorescence images of a *K^LOH^C* metastatic lesion (M #31; TdT, GFP; BF (bright field)). **h**, Pearson correlation matrix relating the number of *Kras^G12D/LOH^* and *Kras^G12D/WT^* metastases, total metastases, tumor width, length and weight, survival, and the percentage of *Kras^G12D/LOH^* area quantified from gross fluorescence imaging of primary tumors. **i**, Number of metastases seeded by *Kras^G12D/WT^*versus *Kras^G12D/LOH^* transplants. N = 22. (*P* = 0.0364). **j**, Pie charts of relative contribution of *Kras^G12D/WT^*(red) and *Kras^G12D/LOH^* (green) cells at metastatic sites: lymph nodes (73% *Kras^G12D/LOH^*), liver (61%), lung (80%) and other sites (75%). **k**, Representative H&E and immunofluorescence (DAPI, TdT, GFP) of a primary PDAC *K^LOH^C* lesion (Scale bar = 200 µm). **l**, Distribution of histological grade by allelic composition of primary tumors (*Kras^G12D/WT^* versus *Kras^G12D/LOH^*; *P* = 0.0351, N = 28 and 17). **m**, Percentage of *Kras^G12D/LOH^* area quantified from gross fluorescence imaging of primary versus metastatic lesions. N = 22. (*P* < 0.0001). Data in (i) and (m) are mean ± s.d. *P* values were calculated using *log-rank (Mantel–Cox) tests* (b), *two-tailed Fisher’s exact tests* (c and l), *χ² test* (e) and *Mann–Whitney U-tests* (i and m). Values in h are *Pearson* correlation coefficients. The diagrams in a, d and f were created using BioRender.

To provide functional validation of this phenomenon and real-time tracing of allelic dynamics in a faithful experimental model of PDAC, we developed a dual reporter system that enables fluorescence based, tissue specific tracing of cancer cells undergoing allele selection at the *Kras* locus. Specifically, we leveraged the unique genomic coordinates of *Kras* and *Rosa26* loci, both residing on murine chromosome 6, ∼30Mb apart, and co-segregating. First, we crossed a *Kras^LSL-G12D^*allele with a *Rosa26^LSL-Cas9-GFP^* to obtain models with the two synthetic cassettes in *cis* (*Kras^LSL-G12D^-Rosa26^LSL-Cas9-GFP^*^)^. Next, we crossed this model to a pancreatic specific *Cre* recombinase allele and an inducible TdTomato (TdT) fluorescent reporter on *Rosa26* in *cis* with the wild-type *Kras* allele to obtain the *K^LOH^C* smGMLT. Tumor initiation is achieved through orthotopic delivery of adeno-associated virus (AAV) particles carrying a combination of single guide RNAs (sgRNAs) targeting the most common tumors suppressor genes in PDAC *Trp53, Cdkn2a* and *Cdkn2b*^15^. The *K^LOH^C* model allows for three potential outcomes: pancreatic cancer cells retaining the wild-type allele (*Kras^G12D/WT^*; TdT+/GFP+), cancer cells with allele selection of the mutant *Kras* and loss of the wild-type (*Kras^G12D/LOH^*; TdT-/GFP+), and tumor cells with allele selection of the wild-type *Kras* and loss of the mutant (*Kras^WT/LOH^*; TdT+/GFP-) (**Figure 1 f, Extended data Figure 2 a-b)**. Performing clinical, gross and histopathological examination of *K^LOH^C* SM-GEMMs, we demonstrated that metastatic lesions are predominantly characterized by *Kras^G12D/LOH^*tumor lineages, and that an increased fraction of the *Kras^G12D/LOH^*population at the primary tumor positively correlates with metastatic burden and poor survival; this phenomenon is not dependent on primary tumor size (P < 0.0001, rho = 0.42) (**Figure 1 g-j, Extended data Figure 2 c-e)**. We did not observe loss of the mutant *Kras^G12D^*allele (*Kras^WT/LOH^*) neither in primary or metastatic lesions, corroborating the concept that mutant *Kras* is a key driver of PDAC progression and it is required for tumor maintainance, thus supporting the notion of oncogene addiction (**Figure 1 k**)^16,17^. This evidence suggests that the timing of the emergence of *Kras^G12D/LOH^*at the primary tumor may be an early indicator of rapid progression to metastatic PDAC **(Extended data Figure 2 f).** Further pathological investigation of *Kras^G12D/WT^*and *Kras^G12D/LOH^* cancer lineages, showed that latter population is associated with high tumor grade and histological transformation to sarcomatoid and anaplastic carcinomas. These phenomena are associated to aggressive disease biology, as previously described by our group (**Figure 1 l-m, Extended data Figure 2 g**)^15,18,19^. Overall these evidences identify *KRAS-*LOH as a critical genomic event associated with of metastatic progression in PDAC.

### *KRAS* LOH is associated with an immunosuppressive phenotype

Functional and genomic data strongly suggest that *KRAS-*LOH contributes to aggressive clinical behavior and metastatic dissemination, potentially supported by phenotypic diversification mirroring mutant allele selection. To corroborate genomic analysis and functional experimental data we perform comprehensive cross-species single cell transcriptomic analysis of publicly available clinical datasets and smGMLT including N = 89 PDAC patients and N = 4 *K^LOH^C* models (**Figure 2 a**).

**Figure 2:**
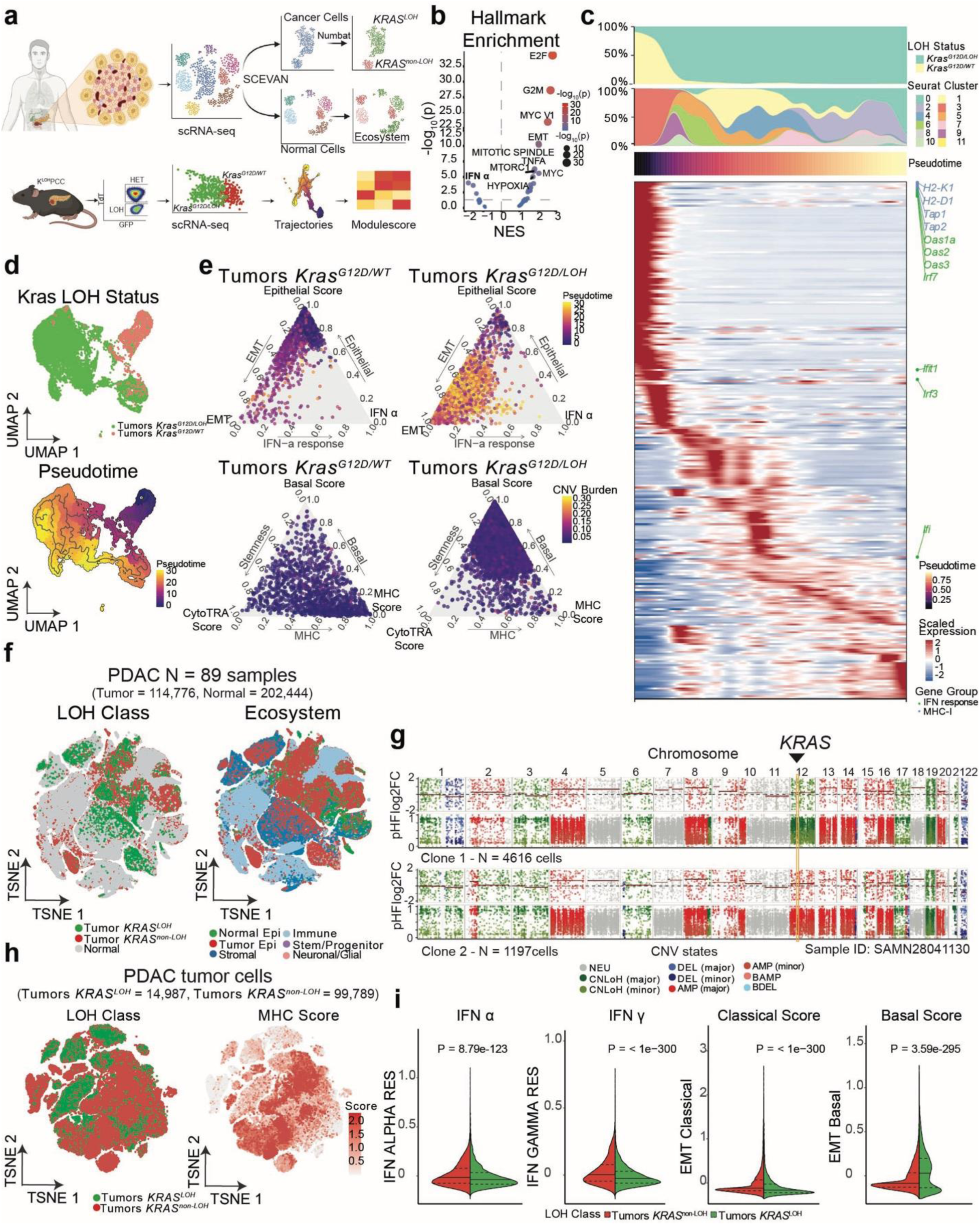
Cross species conserved disruption of interferon response in *KRAS* LOH tumors. **a**, Cross-species scRNA-seq workflow: human PDAC samples (N = 89) were deconvolved (SCEVAN) into cancer cells (N = 126,234), classified as *KRAS^LOH^* or *KRAS^non-LOH^* by Numbat, and normal cells (N = 202,444) annotated to ecosystem; murine *K^LOH^C* tumors were FACS-sorted into *Kras^G12D/WT^* and *Kras^G12D/LOH^* populations by TdT/GFP and profiled by scRNA-seq for trajectory and module-score analysis. **b**, Hallmark pathway enrichment of mouse scRNA sequencing, positive NES values enrichment in *Kras^G12D/LOH^* tumors, negative NES values indicate enrichment in *Kras^G12D/WT^* tumors. **c**, Pseudotemporal ordering of gene programs along the LOH trajectory, annotated by *Kras* LOH status, Seurat cluster and pseudotime, highlighting interferon-response and MHC-I genes. **d**, UMAP of murine cancer cells colored by *Kras* LOH status and pseudotime. **e**, Ternary representations positioning tumor *Kras^G12D/WT^* and *Kras^G12D/LOH^*cells across epithelial, EMT and IFN-α-response programs (top, colored by Monocle3 pseudotime) and across stromal/basal, CytoTRACE score, and MHC score (bottom, colored by CNV burden). **f**, t-SNE of human PDAC cells ( N = 114,776 tumor and N = 202,444 normal cells across 89 samples) labeled by *KRAS* LOH class and ecosystem compartments. **g**, Copy-number inference (per-cell log2 fold change) across chromosomes for two representative malignant clones (Clone 1, N = 4,616 cells; Clone 2, N = 1,197 cells; sample SAMN28041130), with the *KRAS* locus indicated. h, PDAC tumor cells (*KRAS^LOH^*, N = 14,987; *KRAS^non-LOH^*, N = 99,789) colored by LOH class and MHC score. **i**, Hallmark IFN-α, IFN-γ, EMT-classical and EMT-basal module scores in human malignant cells by LOH class (*P* = 8.79 × 10^-^¹²³, *P* = < 1× 10^-300^, *P* = < 1× 10^-300^ and *P* = 3.59 × 10⁻²⁹⁵, respectively). Violin plots show the median and quartiles (25, and 75 quartile). *P* values were calculated using *Wilcoxon test*.

Following fluorescence-based sorting and quality control, we retrieved N = 11,458 *bona fide* cancer cells representative of both *Kras^G12D/LOH^* and *Kras^G12D/WT^* subpopulations derived from the *K^LOH^C* models, as further confirmed by computational analysis of TdTomato transcriptomics (**Extended data Figure 3 a**). Deconvolution of single cell transcriptomes derived from *Kras^G12D/LOH^* and *Kras^G12D/WT^* tumor cells demonstrates that the former population is characterized by the activation of molecular pathways involved in cell proliferation (E2F, G2M, MYC), epithelial-to-mesenchymal transition (EMT), stemness features, and basal-like phenotypes, which have been described by our group and others to characterize aggressive lineages in PDAC (**Figure 2 b-e, Extended data Figure 3 b-f, Methods**) ^8,15,20^. On the other hand, *Kras^G12D/WT^*cells were enriched for pathways characteristic of the classical-like phenotype and involved in interferon (IFN) response, MHC expression, and antigen presentation with significant differential expression of mediators of type I interferon response such as *Irf3* and *Irf7* (**Figure 2 b-e, Extended data Figure 3 g**). Cross species deconvolution of scRNA-seq data from patient-derived PDAC samples provided N = 114,776 malignant and N = 202,444 non-malignant cells respectively (**Figure 2 f, Extended data Figure 4 a-b**). To capture transcriptomic features of *KRAS* LOH, we calculated *KRAS* allelic status through haplotype-aware bioinformatic pipelines (**Figure 2 f-h, Extended data Figure 4 c-d, Methods**). Strikingly, analysis of the transcriptomes of malignant subpopulations with *KRAS* LOH demonstrated a significant suppression of pathways involved in immune activation and interferon response and an upregulation of programs involved in cell plasticity and mesenchymal reprogramming of malignant epithelial tumors including EMT and basal-like signatures, thus mirroring the transcriptomic programs and functional features observed in the smGMLT tumor models (**Figure 2 i, Extended data Figure 4 g-j**). To characterize the tumor microenvironment (TME) remodeling associated with *KRAS* allelic status, we grouped samples based on the fraction of the malignant compartment with *KRAS* LOH (**Methods**). This analysis showed an enrichment for CD8+ T cells with cytotoxic and proliferative features in samples with virtual less than 50% of *KRAS* LOH cells (**Figure 3 a-c, Extended data Figure 5 a-c, Methods**). To further reconstruct the geospatial architecture of PDAC TME as related to the allelic dosage of *KRAS* we performed spatial transcriptomic analysis of pancreatic tumors surgically resected from N = 9 patients (N = 115,104 spots) with a confirmed pathological diagnosis of PDAC and genomic annotation of the *KRAS* allelic status through targeted NGS (**Figure 3 d, Extended data Figure 6 a-d, Methods**). Overall, spatial transcriptomic data deconvolution demonstrated, an enrichment for basal-like features in the malignant compartment and an immunosuppressive microenvironment, characterized by reduced CD8+ T cell activation, transcriptomic signatures of exhausted T cells, and reduced immune-to-tumor cells interactions in *KRAS*-LOH tumors (**Figure 3 e-i, Extended data Figure 6 a-d, Extended data Figure 7 a-e, Methods**). Overall, this cross-species analysis strongly suggests that cancers composed by a high fraction of malignant cells with *KRAS* LOH display an immuno-suppressive phenotype and are characterized by an immune-cold microenvironment.

**Figure 3:**
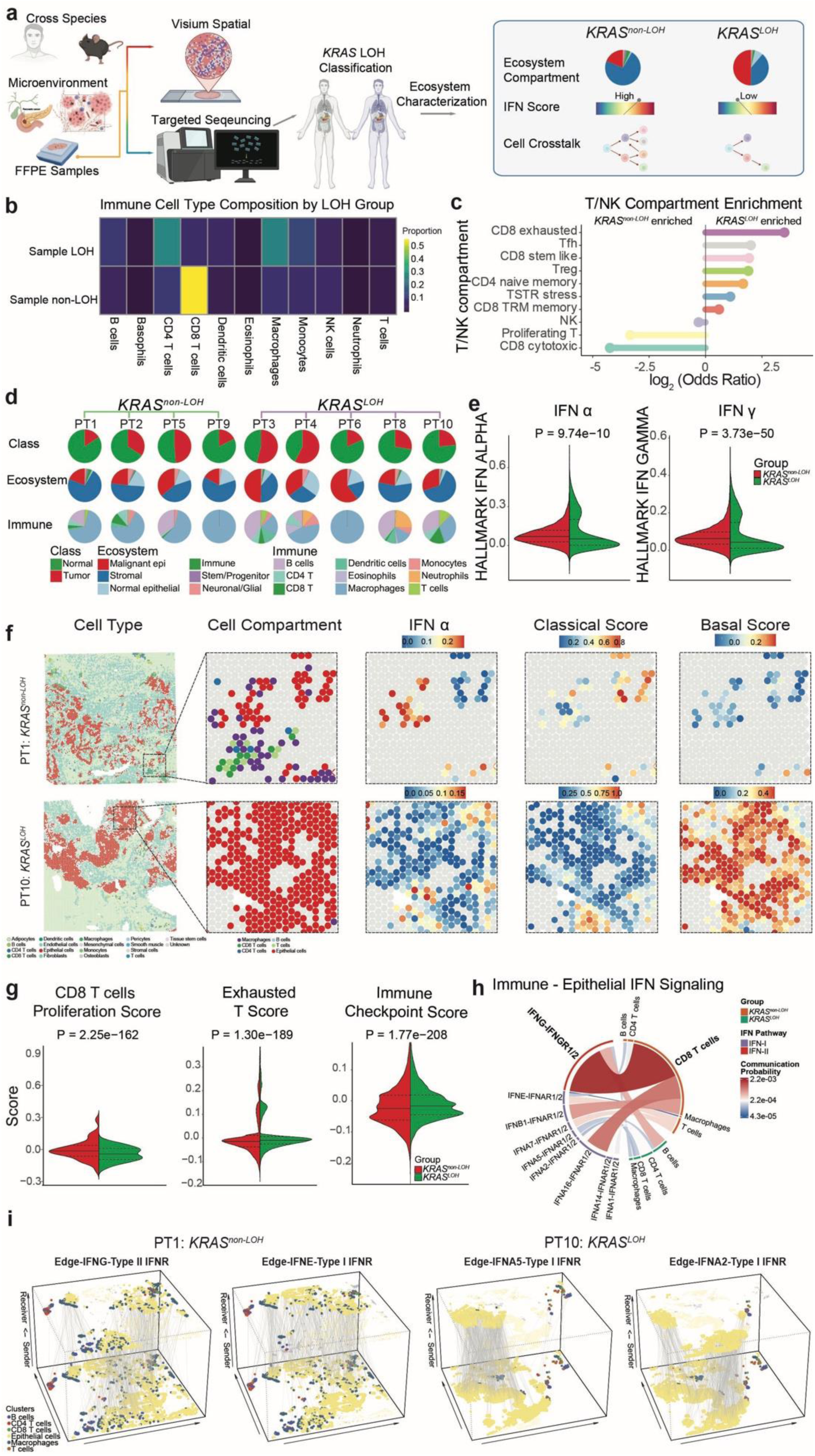
*KRAS* LOH is associated with a CD8-excluded microenvironment in human PDAC. **a**, Cross-species experimental workflow: FFPE human PDAC samples were profiled by Visium spatial transcriptomics and targeted sequencing, classified by *KRAS* LOH status and characterized at the level of the tumor ecosystem, and integrated with the murine data. The inset summarizes the features that distinguish *KRAS^non-LOH^*from *KRAS^LOH^* tumors: ecosystem compartment composition, interferon (IFN) score and cell–cell crosstalk. **b**, Heatmap of immune cell-type composition by sample LOH status. Rows correspond to sample LOH (top, LOH > 50%) and sample non-LOH (bottom, LOH < 50%) and columns to the annotated immune cell types; colour encodes the relative abundance (Proportion) of each cell type within each group, from low (dark) to high (yellow). CD8 T cells are the most strongly enriched population in sample non-LOH group. **c**, Enrichment of T and NK compartments in sample LOH versus sample non-LOH, expressed as the log2∼ odds ratio; positive values denote enrichment in sample LOH (CD8 exhausted, Tfh, CD8 stem-like, Treg, CD4 naive memory, TSTR stress, and CD8 TRM memory) and negative values enrichment in sample non-LOH group (CD8 cytotoxic, proliferating T, and NK). **d**, Per-patient composition of the Class (tumor versus normal), Ecosystem and Immune compartments shown as pie charts for *KRAS^non-LOH^* (PT1, PT2, PT5, PT9) and *KRAS^LOH^* (PT3, PT4, PT6, PT8, PT10) tumors. **e**, Hallmark IFN-α and IFN-γ module scores in malignant cells, compared between *KRAS^non-LOH^*and *KRAS^LOH^* tumors (*P* = 9.74 × 10⁻¹⁰ and *P* = 3.73 × 10⁻⁵⁰, respectively). **f**, Representative spatial maps of a *KRAS^non-LOH^* (PT1) and a *KRAS*^LOH^ (PT10) tumor, showing, from left to right, the spatial distribution of cell types, cell compartment (malignant and immune cells), and IFN-α, classical and basal transcriptional scores on malignant compartment. **g**, CD8 T cell proliferation score, exhausted T cell score and immune-checkpoint score in non-malignant regions, compared between *KRAS^non-LOH^* and *KRAS^LOH^* tumors (*P* = 2.25 × 10⁻¹⁶², *P* = 1.30 × 10⁻¹⁸⁹ and *P* = 1.77 × 10⁻²⁰⁸, respectively). **h**, Chord diagram of inferred immune–epithelial interferon signaling ligand-receptor interactions. The outer sectors indicates cell types, split by LOH group (*KRAS^non-LOH^* and *KRAS^LOH^*); the inner ribbons are individual interferon ligand–receptor pairs (type I, e.g. IFNA2–IFNAR1/2, IFNE–IFNAR1/2; and type II, IFNG–IFNGR1/2), colored by IFN pathway, and ribbon colour intensity encodes communication probability. The IFNG–IFNGR1/2 (type II) interaction towards CD8 T cells dominates in *KRAS^non-LOH^*tumor. **i**, Spatially resolved ligand– receptor signaling edges inferred with CytoSignal for a *KRAS^non-LOH^* (PT1; IFNG–type II IFN receptor, IFNE–type I IFN receptor) and a *KRAS^LOH^* (PT10; IFNA5– and IFNA2–type I IFN receptor) tumor. In each plot the two horizontal axes are the spatial coordinates of the tissue section and points are cells colored by cluster (B cells, CD4 T cells, CD8 T cells, epithelial cells, macrophages, and T cells); grey lines are inferred signaling edges connecting a sender cell (expressing the ligand) to a receiver cell (expressing the cognate receptor), the vertical axis ordering cells from sender to receiver. The density and spatial extent of the edges reflect the magnitude and reach of each interferon interaction within the tissue. In **e** and **g**, violin plots show the median and quartiles. *P* values were calculated using *Wilcoxon test* (**e** and **g**); enrichment in **c** was assessed by *Fisher’s exact test* (*P* < 0.001). The diagram in a was created using BioRender.

### Wild-type *KRAS* is a master regulator of immune-surveillance in pancreatic cancer

Single cell and spatial transcriptomic analysis showed that tumor cells with *KRAS* LOH display an immuno-suppressive phenotype associate with an immune-cold microenvironment, while cells retaining the wild-type allele activate programs related to interferon response and antigen presentation through MHC-I. To test if *KRAS* allelic status differentially regulates immune activation in PDAC, we generated *Kras^G12D/LOH^*and *Kras^G12D/WT^* isogenic tumors from *K^LOH^C* smGMLT and established orthotopic transplants in pancreas of animals with either intact or defective adaptive immunity (C57BL/6J and *Foxn1^nu^*, respectively, **Methods**). Strikingly, tumor growth and penetrance was severly affected in *Kras^G12D/WT^* transplants in C57BL/6J, where only N = 2 *Kras^G12D/WT^* transplants were able to establish tumors upon spontaneous loss of the wild-type allele as demonstrated by the emergence of TdT-/GFP+ *Kras^G12D/LOH^*lesions following long latency, thus suggesting that an intact adaptive immune response exerts a potent selective pressure for the emergence of immune evasive clones characterized by loss of the wild-type allele (**P < 0.0001**, **Figure 4 a, Extended data Figure 8 a-c**). This phenotype was completely abrogated in immunocompromised animals where both isogenic lines established rapidly metastatic tumors (**Figure 4 a, Extended data Figure 8 a-c**). Competition transplantation experiments also confirmed that *Kras^G12D/WT^* cells are unable to participate to tumor initiation and progression, suggesting their counterselection in recipient mice with an intact immune system (**Figure 4 b-c, Extended data Figure 8 d-e**). We therefore postulated that, if the phenomenon here described is *KRAS*-dependent, the reconstitution of wild-type *Kras* allele would be sufficient to restore the interferon signaling response and promote immune recognition *in vivo* in immune competent animals. In line with our hypothesis, reconstitution of the wild-type *Kras* proto-oncogene in *Kras^G12D/LOH^* cells phenocopied *Kras^G12D/WT^* cells upon transplantation, with 90% of the transplants failing to promote primary tumor and metastatic growth *in vivo* (**Figure 4 c-d, Extended data Figure 8 f**). Transcriptomic and cytofluorimetric analysis of wild-type *Kras* reconstituted *Kras^G12D/LOH^*cells showed that wild-type allele is sufficient to activate signatures related to an interferon response, including expression of MHC-I (**Figure 4 e-g, Extended data Figure 9 a-d**). These functional experiments strongly support the hypothesis that the *KRAS* proto-oncogene is a master modulator of tumor immune response and acts as a tumor suppressor gene in the context of metastatic pancreatic cancer.

**Figure 4:**
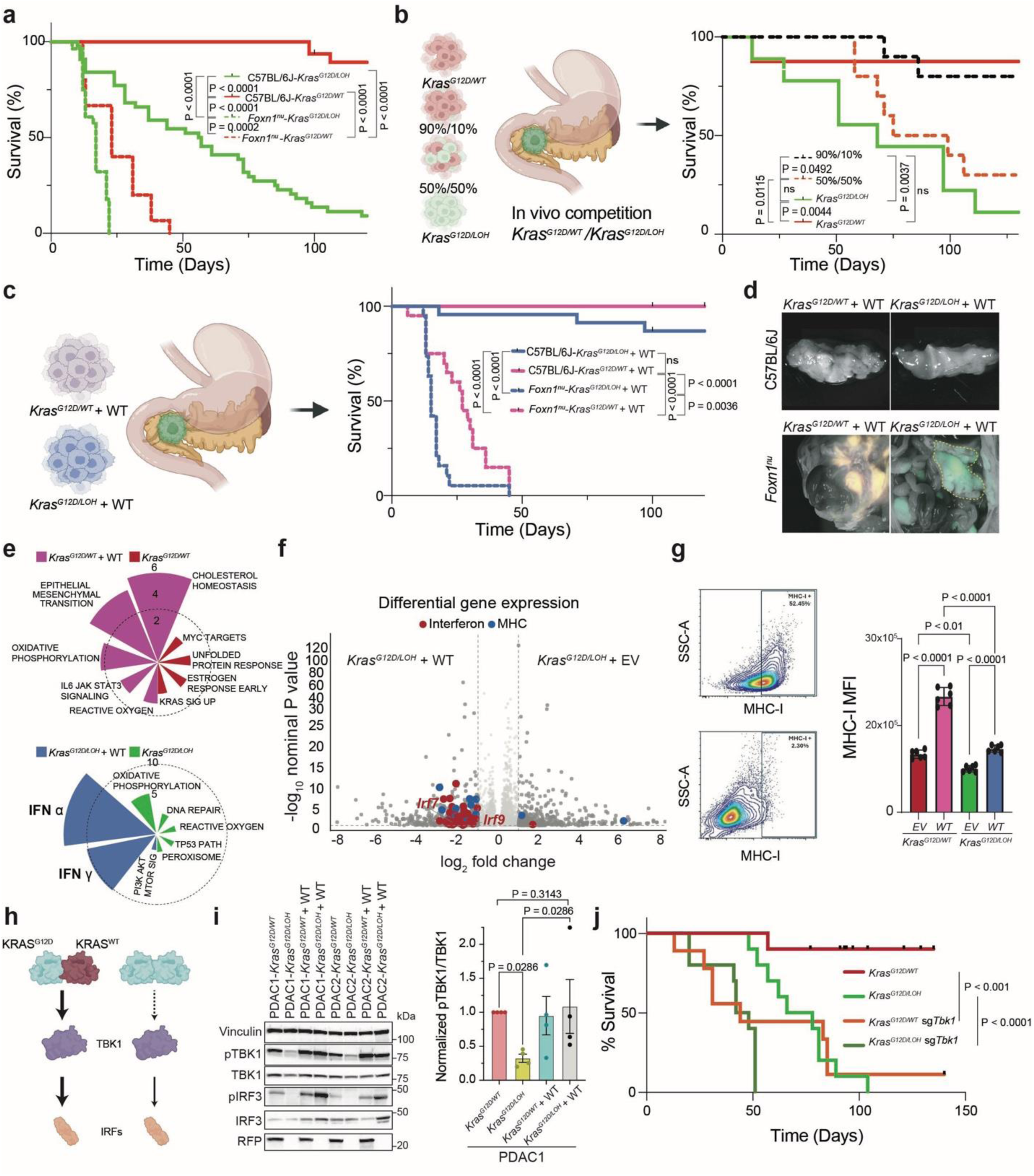
Wild-type *KRAS* mediates a potent anti-tumor immune response. a, Survival curves of immunocompetent C57BL/6J and *Foxn1^nu^*recipients after orthotopic transplantation of sorted *Kras^G12D/LOH^* or *Kras^G12D/WT^* cells isolated from *K^LOH^C* primary tumors. **b**, Competitive orthotopic transplantation survival curves of the two populations within the same tumor: *Kras^G12D/WT^*and *Kras^G12D/LOH^* cells were mixed at 90:10 and 50:50 ratios (schematic, left) and transplanted into immunocompetent recipients, with survival compared to mice receiving each population alone (right). **c**, Survival curves of recipients transplanted with wild-type-reconstituted cells. Sorted *Kras^G12D/LOH^* and *Kras^G12D/WT^* cells in which the wild-type *Kras* allele was reconstituted (+WT) or transduced with empty vector as control. **d**, Representative of N = 77 fluorescence microscopy images of the pancreas from the four arms of the experiment in **c**. **e**, Gene set enrichment analysis of sorted populations, shown as radial plots of enriched Hallmark pathways: *Kras^G12D/WT^* versus *Kras^G12D/WT^* + WT (top) and *Kras^G12D/LOH^* versus *Kras^G12D/LOH^* + WT (bottom). Bar length reflects the magnitude of enrichment and colors the direction of regulation. **f**, Volcano plot of differential gene expression between *Kras^G12D/LOH^* + EV (empty-vector control) and *Kras^G12D/LOH^* + WT cells; interferon-stimulated genes (red; e.g. *Irf7*, *Irf9*) and MHC-I genes (blue; *H2-Q4*, *H2-Q6*, *H2-T23*, *H2-K1*, *H2-D1*, *H2-Q10*) are enriched in the wild-type-reconstituted cells. **g**, Flow-cytometric quantification of cell-surface MHC-I: representative gating of MHC-I⁺ cells against forward scatter (FSC-A) (left) and MHC-I mean fluorescence intensity (MFI) for empty-vector (+EV) and wild-type-reconstituted (+WT) *Kras^G12D/WT^* and *Kras^G12D/LOH^* cells (right). **h**, Schematic showing proposed differential activation of downstream TBK1-IRF axis in presence or absence of wild-type *KRAS*. **i**, Immunoblot and quantification of phospho-TBK1 (pTBK1) and total TBK1, phospho-IRF3 (pIRF3) and total IRF3, with vinculin as loading control and TdT (RFP) as a lineage marker, across two independent PDAC lines (PDAC1, PDAC2) of the indicated genotypes (*Kras^G12D/WT^*, *Kras^G12D/LOH^*, *Kras^G12D/WT^* + WT, *Kras^G12D/LOH^* + WT); IRF3 phosphorylation tracks with the presence of the wild-type allele. Data in i are mean ± s.e.m. *P* value was calculated using the *Mann–Whitney test* (N = 4). **j**, Kaplan–Meier survival of immunocompetent recipients transplanted with *Kras^G12D/WT^*, *Kras^G12D/LOH^*, *Kras^G12D/WT^*;sg*Tbk1* and *Kras^G12D/LOH^*;sg*Tbk1* cells; CRISPR ablation of *Tbk1* shortens survival in both genotypes, consistent with a tumor-restraining function of the axis. Data in **g** are mean ± s.d. *P* values were calculated using log-rank (*Mantel– Cox) tests* (**a**, **b**, **c** and **j**) and *one-way ANOVA* with multiple-comparison correction (**g**); *P* values are as indicated in each panel. The diagrams in **b**, **c** and **h** were created using BioRender.

To provide mechanistic understanding of the phenotype observed in transplantation studies, we dissected canonical downstream signaling activated by the *Kras* oncogene in *Kras^G12D/LOH^* and *Kras^G12D/WT^*cells (**Figure 4 h**). Strikingly, we observed a major difference in the engagement of the KRAS-TBK1-pIRF3 axis between *Kras^G12D/LOH^* and *Kras^G12D/WT^* cells (**Figure 4 i**). Similarly, reconstitution of wild-type *Kras* in *Kras^G12D/LOH^* cells resulted in a basal upregulation of pTBK1 (**Figure 4 i**). Thus, wild-type Kras is sufficient to differentially activate TBK1 and pIRF3. To demonstrate that the KRAS-TBK1-IRF3 pathway is a major contributor of the phenotype, we performed loss-of-function studies and transplantation experiments which demonstrated that ablation of *Tbk1* partially phenocopies loss of the wild-type allele in *Kras^G12D/WT^* cells and promote immune evasion and metastatic progression (**Figure 4 j**). Concluding, this experimental evidence strongly suggests that wild-type KRAS differentially participate to TBK1 activation when compared to mutant KRAS.

## Discussion

Here we provide compelling functional evidence for the role of *KRAS* allele selection during pancreatic cancer progression. Allelic imbalance of oncogenic drivers is a frequent feature of solid tumor evolution. While the field has focused on the gain of mutant dosage as a critical genomic driver of progression, we revise this framework and provide evidence that the wild-type allele is not a passive bystander of tumor evolution, but an actively counter-selected oncosuppressor.

Our results underscore the importance of tracing genomic evolution of cancer driver mutations to define hallmark events in clinical progression. The smGMLT is complementary to lineage-tracing systems that report allelic loss at a driver loci, and CRISPR-based lineage recorders that reconstruct clonal dynamics during metastatic dissemination^21–23^. We argue that the continued development of such platforms is critical to resolve how individual driver events shape phenotypic transitions and the metastatic process.

Contextualizing our results lead us to the broader concept that allelic imbalance of oncogenic drivers plausibly follow tissue-restricted evolutionary trajectories, shaped by signaling pathway dosage, dependency, differentiation state, and microenvironmental cues. Such a model could explain the observed difference in frequency of *KRAS* LOH across tumor types. If PDAC dominates the landscape of proto-oncogene loss, this may reflect not only a greater tolerance for oncogenic activation and a greater pathway addiction as previously postulated, but also the presence of critical microenvironmental restrains that substantially contribute to the evolutionary bottlenecks shaping the clonal landscape and the evolutionary routes to metastasis. Other *KRAS*-driven malignancies, such as lung adenocarcinoma and colorectal carcinoma, potentially leverage alternative immune-evasive evolutionary routes, converging on phenotypes while diverging in genotypes. Indeed, epithelial cancers recurrently converge over canonical phenotypic trajectories, including de-differentiation and lineage infidelity, acquisition of stem-like properties, and mechanisms to support immune-evasion^15,24–26^. Recent work leveraging mouse models and lineage tracing barcode technologies have shown that high-plasticity states are responsible for lung cancer progression and metastasis, while a basal-like phenotype converge on the activation of an immuno-suppressive response in PDAC^20,24,25^.

Taken together, our work brings forward a counterintuitive evolutionary mechanism for the selection of mutant *KRAS*, strongly implying that wild-type *KRAS* functions as a potent oncosuppressor in advanced disease by enabling immune evasion. Previous studies have described how wild-type *KRAS* can function as a cell-intrinsic oncosuppressor, reducing MAPK dosage through dimerization with the mutant protein^27^. Conversely, oncogenic KRAS engages an RSK1/NF1 feedback loop that selectively suppresses wild-type RAS signaling. Our data extend this principle into a cell-extrinsic, immune-mediated phenotype that acts as a driver of the positive selection of *KRAS* LOH during systemic dissemination. Consistently, we observed a coordinated reduction of the interferon-response and MHC-I expression, a configuration expected to compromise antigen presentation and blunt intra-tumor CD8⁺ T cell recognition.

This concept lends mechanistic weight to the long-standing clinical observation of an immune-cold microenvironment in PDAC, and partially offers a genetic and evolutionary explanation whereby one discrete genomic event fuels addiction to oncogenic and anabolic pathways (MAPK, PI3K-AKT) and disrupts a critical tumor suppressive function, promoting immune evasion and metastatic spread. However, only a fraction of human PDAC exhibit loss of the wild-type *KRAS* allele, suggesting that other mechanisms may be alternative to proto-oncogene loss to drive immune evasion^2,28^. For example, clinical-genomic studies have shown that the loss of IFN ligands on chromosome 9p may be a critical evolutionary event to drive immune evasion in several cancer types including pancreatic cancer^29^. On the same topic, our group has shown that loss of chromosome 21q, harbouring critical IFN receptors, is sufficient to drive aggressive metastatic behavior in kidney cancers by downmodulating an IFN receptor mediated response^30^. Whether these routes are mutually exclusive or complementary remains an open and clinically relevant question.

Our findings also raise the possibility that *KRAS* allelic status could serve as a surrogate for a defined immune-evasive state. To support this proposition, several groups have observed that treatment with mutant-selective KRAS inhibitors are able to evoke a potent immunological response in pre-clinical models, thus supporting the finding that pharmacological pressure to retain the wild-type copy may sustain a beneficial interferon mediated response^31–35^. Activation of interferon signaling in response to KRAS inhibitors has also been recently observed in colorectal cancer^36^. The authors propose that the activation of the interferon response emerges early during treatment with KRASi and it is partially regulated by TBK1. Recent work on a large cohort of PDAC organoids showed that downregulation of the interferon signaling is associated with resistance to KRAS inhibitors^37^. In line with our experimental data, we propose that this phenomenon is mediated by the positive selection of the proto-oncogene under the pharmacological pressure of mutant-selective KRAS inhibitors. Altogether, these observation further highlight the concept of tumor and tissue specificity of oncogene addiction in the context of the evolutionary dynamics of a given cancer type.

The differential biochemical activation of the downstream effectors of KRAS is of critical importance in the era of KRAS inhibitors. We propose that the a wild-type copy of *KRAS* is, indeed, sufficient to promote the activation of the TBK1-IRF3 pathway. We also speculate that targeting KRAS would be an amenable pharmacological strategy to treat tumors retaining the proto-oncogene; however, whether the KRAS-LOH status have a clinically meaningful effect in selecting patient for pan-RAS versus mutant selective inhibitors has yet to be proven and a topic for future investigations. Overall, evidence obtained by our group and others strongly suggests that the intrinsic instability of the cancer genomes is a key driver of tumor progression^15,30,38^. The cooperation between ongoing chromosomal instability and allele selectionat the *KRAS* locus may pose a critical selective pressure to reduce the tone of activation of TBK1, evoked by both KRAS and cGAS-STING signaling^39,40^. We therefore conclude that the counter-selection of the proto-oncogene results in multiple benefits for cancer cells that ultimately converge on an chromosomally unstable, de-differentiated, immune-evasive phenotype.

## Methods

### Genetic engineered mouse models

The *Rosa26^LSL-Cas9-GFP^* strain was generated in F. Zhang’s laboratory and obtained through the Jackson Laboratory (024857)^41^. The *Rosa26^LSL-TdT^* strain was generated in H. Zeng’s laboratory and obtained through the Jackson Laboratory (007908)^42^. The *Kras^LSL-G12D^* strain was generated in T. Jacks laboratory and obtained through the Jackson Laboratory (008179)^43^. The *Tg(Pdx1-Cre)* strain was generated by D. Tuveson and obtained through the Jackson Laboratory (014647)^6^. The *K^LOH^C (Tg(Pdx1-Cre)^LSL^-Kras^G12D/WT^Rosa26^LSL-TdT/LSL-Cas9-GFP^)* mouse model was generated through a multistep breeding strategy. First, mice carrying the *Kras^LSL-G12D^* allele were crossed with *Rosa26^LSL-Cas9-GFP^* mice to obtain mice carrying the two synthetic cassettes in *cis*. In parallel, mice carrying the *Rosa26^LSL-TdT^* allele were crossed with mice carrying the pancreatic-specific *Tg(Pdx1-Cre)* recombinase allele. Mice carrying *Kras^LSL-G12D^* and *Rosa26^LSL-Cas9-GFP^*were subsequently crossed with mice carrying *Tg(Pdx1-Cre)* and *Rosa26^LSL-TdT^* to generate the *K^LOH^C* mouse model. The *Rosa26* locus is located on murine chromosome 6, approximately ∼30 Mb from the *Kras* locus. In the *K^LOH^C* model, the *Rosa26^LSL-Cas9-GFP^* and *Kras^LSL-G12D^* alleles are present in *cis*, whereas the *Rosa26^LSL-TdT^*reporter is located in *trans* with the *Kras^LSL-G12D^* allele. Strains were kept in a C57BL/6J background. All animal studies and procedures were approved by the UT MD Anderson Cancer Center Institutional Animal Care and Use Committee. All experiments conformed to the relevant regulatory standards and were overseen by the institutional review board. For orthotopic tumors, mice were euthanized on symptoms of distress or with bulky disease. No sex bias was introduced during the generation of experimental cohorts. Littermates of the same sex were assigned randomly to experimental arms.

### GEMM nomenclature

The following GEMM nomenclature was used: *K^LOH^C, (Tg(Pdx1-Cre)^LSL^-Kras^G12D/WT^Rosa26-^LSL-TdT/LSL-Cas9-GFP^)*. *K^LOH^C* model was tested for the following combination of AAV-cre: *sgTrp53, sgCdkn2a and sgCdkn2b*^15^.

### Tumor cell isolation, flow cytometry, cell sorting and culture

*Ex vivo* cultures from primary tumor explants were generated by mechanical dissociation and incubation for 1 h at 37 °C with a solution of collagenase IV/dispase (2 mg ml−1) (Invitrogen), resuspended in DMEM (Lonza) and filtered. Cells derived from tumor dissociation and digestion were resuspended as single-cell suspensions in 1× PBS and 5% FBS. Single-cell suspensions were processed using fluorescence-activated cell sorting (FACS) using GFP fluorescence as a reporter for tumor cells and TdTomato fluorescence as a reporter of LOH (*Kras^G12D/WT^*; TdT+/GFP+ and *Kras^G12D/LOH^*; TdT-/GFP+). Single-cell suspensions were sorted using the BD Aria II cell sorter. Sorted GFP+ and TdT+/-cells were centrifuged at 200g for 5 min and resuspended in the appropriate medium. Tumor explans were plated at high confluency in 96-well plates with DMEM/F12 supplemented with 1% penicillin–streptomycin, 10% FBS and incubated with AAV (10^9^ viral particles) for 8 h at 37 °C 5% CO2. Single-cell suspensions were collected, pelleted and embedded in Matrigel or transplanted, as previously described^15,18^.

### Animal procedures

#### Orthotopic parenchymal pancreatic injection

Tumor-explant-derived two cultures were resuspended in single-cell suspension using trypsin-based method at a concentration of 5,000 cells per µl in 1:1 mixture of Matrigel and 1× PBS. Mice were anaesthetized using isofluorane (Henry Schein Animal Health). Shaved skin was disinfected with betadine and 70% ethanol and 1 cm incisions were performed through the skin/subcutaneous and muscular/peritoneal layers. The spleen and tail of the pancreas were identified and exposed, and one injection was performed in the pancreatic tail and body. The muscular/peritoneal planes were closed using continuous absorbable sutures. The skin/subcutaneous planes were closed using interrupted absorbable sutures. Analgesia was achieved with Ethiqa XR (3.25 mg per kg, once). *Euthanasia, necropsy and tissue collection*

Mice were euthanized by exposure to CO2 followed by cervical dislocation. tumor size and weight, infiltrated organs, and metastasis number and location were annotated. Fluorescence and bright-field images were acquired using the Leica system. Euthanasia was performed with animals at clinical terminal disease and metastatic tumor burden.

### Recombinant DNA

Packages of gRNAs were designed and synthetized according to the following scheme: EcorI restriction site – U6 promoter – gRNA1 sequence – gRNA scaffold – polyA – U6 promoter – gRNAn sequence – gRNA scaffold – polyA – AscI restriction site. The synthetic sequence was assembled into the pEMS2158 AAV vector (Addgene no. 70119) into the EcorI and AscI restriction sites. The following plasmids were used in this study: pHAGE-KRAS is a gift from Drs. Mills and Scott and validated elsewhere (Addgene plasmid # 116755)^44^; pHAGE-puro is a gift from Dr. Vacok and validated elsewhere (Addgene plasmid # 118692)^45^.

### sgRNA design

The following sgRNA sequences used in this study were obtained through GenScript: *Trp53*, CATAAGGTACCACCACGCTG; *Cdkn2a*, TATAATTATTGGGTCAGGGG; *Cdkn2b*, GTCATGAGACAAATAGGCGG; *Tbk1*, TTTGAACATCCACTGGGCGA.

### Virus production

Plasmid DNA preparations were generated using endotoxin-free MIDI kits (Qiagen). Large-scale AAV particle production was outsourced to Vector Biolab (1013 IU ml−1).

Lentiviral particles were produced as previously described^46^. In brief, infectious viral particles were produced using psPAX2 and pMD2G helper plasmids. For transfection, HEK293T cells were cultured in DMEM containing 10% FBS (Gibco), 100 IU ml−1 penicillin (Gibco), 100 μg ml−1 streptomycin (Gibco) and 4 mM caffeine (Sigma-Aldrich) and transfected using the polyethylenimine method. Virus-containing supernatant was collected 48–72 h after transfection, centrifuged at 3,000 rpm for 10 min and filtered through 0.45 μm low-protein-binding filters (Corning). High-titer preparations were obtained by multiple rounds of ultracentrifugation at 23,000 rpm for 3 h each.

### Immunohistochemistry and immunofluorescence

Tumor samples were fixed in 4% formaldehyde for 24 h at room temperature, moved into 70% ethanol for 12 h, and then embedded in paraffin (Leica ASP300S). After cutting (Leica RM2235) and baking at 60 °C for 20 min for de-paraffinization, slides were treated with Citra-Plus Solution (BioGenex) for antigen unmasking according to the manufacturer’s instructions. For immunohistochemical staining, endogenous peroxidases were inactivated by 3% hydrogen peroxide at room temperature for 15 min. Non-specific signals were blocked using 5% BSA and 5% goat serum for 1 h. Tumor samples were stained with primary antibodies for 12 h at 4 °C and the Mouse on Mouse Kit (Vector Laboratory) was used when appropriate according to the manufacturer’s instructions. Antibodies used were as follows: RFP (Thermo Fisher Scientific, MA5-15257, 1:200), GFP (Abcam, 13970, 1:200), KRT19 (Abcam, 52625, 1:100), vimentin (Abcam, 137321, 1:100). For immunostaining, ImmPress (Vector Laboratory) were used as secondary antibodies and Nova RED (Vector Laboratory) was used for detection. Images were captured with a Nikon DS-Fi1 digital camera using a wide-field Nikon Eclipse Ci microscope. For immunofluorescence, secondary antibodies conjugated to Alexa488, Alexa647 and Alexa555 (Molecular Probes) were used. Images were captured with a Hamamatsu C11440 digital camera, using a wide-field Nikon EclipseNi microscope and a Nikon high-speed multi-photon confocal microscope A1 R MP. Total staining score was weighed to the intensity and prevalence (percentage of positive tumor cells and intensity score of 0 to 3) in random fields at 20× magnification. Quantitative analysis was performed using Image J and Immunoratio programs according to the providers’ instructions at 20× original magnification.

### Protein extraction and western blot analyses

For cellular protein lysates, cells were scraped on ice using cold Ripa lysis buffer (150 nM NaCl, 50 mM Tris HCl pH 8, 1% Igepal, 0.5% sodium deoxycholate, 0.1% SDS) supplemented with a HALT protease and phosphatase inhibitor cocktail (ThermoFisher). Cell lysates were centrifuged at 17,500*g* for 20 min at 4 °C and supernatants were collected. Proteins were separated in 4–20% SDS–PAGE (Criterion Precast Midi Gel, Bio-Rad) and transferred to nitrocellulose membranes (Trans-Blot Turbo Midi 0.2-μm nitrocellulose transfer pack, Bio-Rad). Membranes were blocked with 5% nonfat dried milk in PBS and incubated at 4 °C overnight with primary antibodies (RFP: ab62341 Abcam, dilution 1:1000; Vinculin #4650 Cell Signaling Technology, dilution 1:1000; TBK1 #3504 Cell Signaling Technology, dilution 1: 1000; pTBK1 #5483 Cell Signaling Technology, dilution 1:1000; IRF3 #4302 Cell Signaling Technology, dilution 1:1000; pIRF3 #4947 Cell Signaling Technology, dilution 1:1000). Membranes were washed in PBS and incubated for 1 h at room temperature with the appropriate horseradish peroxidase-conjugated secondary antibodies (Cell Signaling Technology) for ECL (enhanced chemoluminescence) detection (SuperSignal WEST Pico PLUS Chemiluminescent Substrate, ThermoFisher).

### Murine scRNA-seq of *Kras^G12D/WT^* and *Kras^G12D/LOH^*

Murine scRNA-seq was performed as previously described^15^. In brief, scRNA-seq was performed on the Chromium platform (10x Genomics) using the Chromium Single Cell 3′ Reagent Kits v3 kit manual version CG000183 Rev C (10x Genomics). The final libraries were loaded onto the NovaSeq 6000 platform (Illumina) to obtain 50,000 reads per cell.

### Processing of murine scRNA-seq data of *Kras^G12D/WT^* and *Kras^G12D/LOH^*

Raw scRNA sequencing fastq files were pre-processed and aligned to the mm10 reference using CellRanger Single Cell Software Suite (v.7.1.0, 10X Genomics)^47^. Count matrices were imported into Seurat (v5.5.1) for quality control and clustering^48^. Doublets were identified with scDblFinder R package (v1.21.2) using the default parameters^49^. After removal doublets, cells with more than 25% counts assigned to mitochondrial genes and cells with less than 500 genes by counts were removed from the analysis. Harmony (v1.2.3) was used for batch-effect correction (https://github.com/immunogenomics/harmony?tab=readme-ov-file). To include only tumor cells in our analysis, we used SCEVAN (v1.0.1) (https://github.com/AntonioDeFalco/SCEVAN) to retrieve malignant cells based on inferred copy-number profiles. Module scores were calculated using Seurat’s AddModuleScore function for Hallmark interferon alpha (v2025.1.Mm), Hallmark interferon gamma (v2025.1.Mm), Hallmark G2M checkpoint (v2026.1.Mm), Hallmark MYC targets V1 (v2026.1.Mm), Hallmark MYC targets V2 (v2026.1.Mm), and Hallmark epithelial mesenchymal transition (v2025.1.Mm.gene) gene sets, as well as EMT Classical, EMT Basal, and Thierry EMT gene sets. The MHC score was similarly calculated using *H2-K1, H2-D1, B2m, Tap1, Tap2*. Cell cycle information was assigned using Seurat’s CellCycleScoring function. Cell type information was inferred by SingleR (v2.8.0) (https://github.com/dviraran/SingleR). CytoTRACE2 (v1.1.0) was applied to predict cellular potency and absolute developmental potential ^50^. Monocle3 (v1.4.29) was used for trajectories analysis ^51^. *Kras* LOH information was inferred based on TdTomato expression at single cell level, with the TdTomato-negative (TdT-) expression population representing cells with *Kras^G12D/LOH^*, and TdTomato-positive (TdT+) expression population representing cells with *Kras^G12D/WT^*. Gene Set Enrichment (GSEA) was performed using the fgsea R package (v1.38.0) by applying gene sets from the Molecular Signature Database (MSigDB, Mus Musculus) via msigdbr (v.26.1.1) R package. Ranked gene lists generated from the DESeq2 analysis were used and normalized enrichment scores (NES) were calculated to identify pathways enriched in *Kras^G12D/LOH^* or *Kras^G12D/WT^* tumor cells. Positive NES values enrichment in *Kras^G12D/LOH^*tumors, whereas negative NES values indicate enrichment in *Kras^G12D/WT^*tumors.

### Processing of human scRNA-seq data of *KRAS^non-LOH^* and *KRAS^LOH^*

Publicly available human PDAC scRNA-seq data from (PRJNA647411, PRJNA657844, PRJNA664962, PRJNA806978, PRJNA834096) GSE154778, GSE211644, GSE158356, GSE194247, GSE202051, and GSE156405 were downloaded from NCBI SRA. Raw scRNA-seq fastq files were pre-processed and aligned to human reference GRCh38 using CellRanger Single Cell Software Suite (v7.1.0, 10X Genomics), Count matrices were imported into Seurat (v.5.5.1) for quality control and clustering^48^. Doublets were identified with scDblFinder (v1.21.2) using the default parameters^49^. After removal doublets, cells with more than 25% counts assigned to mitochondrial genes and cells with less than 500 genes by counts were removed from the analysis. N = 114,776 malignant and N = 202,444 non-malignant cells respectively collected from N = 89 independent lesions were included for the final analysis. Harmony (v1.2.3) was used for batch-effect correction (https://github.com/immunogenomics/harmony?tab=readme-ov-file). To include only tumor cells in our analysis, we used SCEVAN (v1.0.1) (https://github.com/AntonioDeFalco/SCEVAN) to retrieve malignant cells based on inferred copy-number information. Numbat (v1.4.2)^52^ was applied to infer haplotype information across the genome, genomic regions with a copy number state annotated as LOH were subset and mapped to the *KRAS* genomic region (hg38, chr12:25205246-25250929), cells with *KRAS* region annotated as LOH were assigned to *KRAS^LOH^* and cells without *KRAS* region called LOH were assigned to *KRAS^non-LOH^*. Module scores were calculated using Seurat’s AddModuleScore function for Hallmark interferon alpha (v2025.1.Hs), Hallmark interferon gamma (v2025.1.Hs) gene sets, as well as EMT Classical, EMT Basal genes, and Thierry EMT gene sets. The MHC score was similarly calculated using *HLA-A, HLA-B, HLA-C, B2M, TAP1, TAP2*. Cell cycle information was assigned using Seurat’s CellCycleScoring function and cell types were annotated by SingleR (v2.8.0) (https://github.com/dviraran/SingleR). Gene Set Enrichment (GSEA) was performed using the fgsea (v1.38.0) R package with gene sets from the Molecular Signature Database (MSigDB, Homo sapiens) via msigdbr (v.26.1.1) R package. Ranked gene lists generated from the DESeq2 analysis were used and then normalized enrichment scores (NES) were calculated to identify pathways enriched in *KRAS^LOH^* or *KRAS^non-LOH^*tumors. Positive NES values enrichment in *KRAS^LOH^* tumors, whereas negative NES values indicate enrichment in *KRAS^non-LOH^* tumors.

### T/NK cell state signature scoring and compartment enrichment analysis of PDAC Human scRNA Seq

T/NK cell states were assigned in RStudio (v2026.9.0.174) using Seurat’s (v5.5.1) AddModuleScore function. Gene sets representing major T-and NK-cell functional states were defined as follows: CD8 cytotoxic (*CD8A, CD8B, GZMB, GZMH, PRF1, NKG7, GNLY*, and *IFNG*), CD8 exhausted (*PDCD1, HAVCR2, LAG3, TIGIT, CTLA4, TOX, CXCL13*, and *ENTPD1*), CD8 stem-like (*TCF7, LEF1, CD28, IL7R, CCR7, SELL, BCL2, ID3, SATB1*, and *CXCR6*), CD8 tissue-resident memory (TRM)/memory (*CD8A, CD8B, ITGAE, CD69, CXCR6, GZMK*, and *IL7R*), CD4 naïve/memory (*CD4, IL7R, CCR7, TCF7, LEF1, LTB*, and *SELL*), regulatory T cell (Treg; *FOXP3, IL2RA, CTLA4, IKZF2, TIGIT, TNFRSF18*, and *TNFRSF4*), T follicular helper (Tfh; *CXCR5, PDCD1, ICOS, BCL6, TOX2, IL21*, and *MAF*), proliferating T cell (*MKI67, TOP2A, STMN1, UBE2C*, and *HMGB2*), T-cell stress response (TSTR)/stress (*HSPA1A, HSPA1B, HSPH1, DNAJB1, HSP90AA1, FOS*, and *JUN*), and NK cell (*NKG7, GNLY, KLRD1, KLRF1, FCGR3A, TYROBP*, and *PRF1*) signatures. Module scores were calculated for each cell based on the average expression of genes within each signature, and T/NK cell state were assigned based on the highest score.

T/NK-cell compartments composition were deconvoluted according to sample *KRAS^LOH^* status (Samples with <50% *KRAS^LOH^* tumor cells were grouped into *KRAS^non-LOH^* and Samples with ≥50% *KRAS^LOH^* tumor cells were grouped into *KRAS^LOH^*), the enrichment of each T/NK compartment was compared between *KRAS^LOH^* and *KRAS^non-LOH^* samples using *Fisher’s exact test*. The direction of enrichment was determined by comparing the proportion of each compartment among T/NK compartments between the *KRAS^LOH^* and *KRAS^non-LOH^* samples and the higher proportion was designated as the enriched group.

### Collection of samples from patients and Visium libraries preparation

Formalin-fixed paraffin-embedded (FFPE) tissue section from patients (N = 9) diagnosed of PDAC were included in this study. Transcriptomic libraries were prepared following the 1.1 x 1.1 mm Visium CytAssist Spatial Gene Expression protocol. H&E-stained images were scanned at 40× using the Vectra Polaris Automated Quantitative Pathology Imaging System (Akoya Biosciences) at 40× resolution; Visium slide alignment was performed using the CytAssist instrument (software version 2.0.1.6). The constructed libraries were pooled at a multiplex level and sequenced on the NovaSeq X Plus with 10B flow cells (Illumina) to achieve a minimum sequencing depth of 25,000 read pairs per tissue-covered spot. *KRAS* allele grouping was based on targeted sequencing results.

### Processing of targeted sequencing data of human FFPE samples

Genomic DNA was quantified using the Equalbit™ 1× dsDNA HS Assay Kit (Vazyme Biotech Co., Nanjing, China), and DNA integrity was assessed by agarose gel electrophoresis. Targeted sequencing was performed using the CleanPlex^®^ OncoZoom^®^ Cancer Hotspot Panel kit (Paragon Genomics, Hayward, CA, USA) according to the manufacturer’s instructions. This multiplex PCR-based targeted resequencing assay is designed for rapid detection of somatic mutations across the hotspot regions of 65 oncogenes and tumor suppressor genes, covering over 2,900 known cancer-associated hotspots. Briefly, 20 ng of input DNA was used for library preparation. Target-specific libraries were generated using the CleanPlex^®^ workflow, which employs multiplex PCR amplification to enrich genomic regions of interest. Following amplification, PCR products underwent enzymatic background cleaning to remove non-specific products, followed by a second PCR to incorporate sample-specific indexing primers, enabling multiplexed sequencing.

The resulting libraries were purified using magnetic beads, and library quality and quantity were assessed with Qubit 2.0 DNA HS Assay (ThermoFisher, Massachusetts, USA), Tapestation High Sensitivity D1000 Assay (Agilent Technologies, California, USA), and QuantStudio^®^ 5 System (Applied Biosystems, California, USA). Equimolar pooling of indexed libraries was performed based on the QC values, and the pooled library was sequenced on the Illumina^®^ NovaSeq™ X Plus platform (Illumina, California, USA) using a paired-end 150 bp (PE150) read configuration, targeting approximately 3 million paired-end reads per sample (i.e., 1.5 million reads per direction).

Raw sequencing fasta files were applied to FastQC (v0.12.1) for quality controls, adapters were trimmed by Trimmomatic (v0.40)^53^. Reads were aligned to human genome reference (GRCh38) using Burrows-Wheeler Aligner (v0.7.19) with default settings^54^. BAM files were further processed according to GATK best practices, including mark of duplicated reads, realignment around indels and base recalibration using GATK (v4.4.0.0)^55^. Somatic mutations were called by Mutect2^56^. All identified mutations were filtered for sequencing depth (coverage > 30X) and only variants with a “PASS” filter status were included in subsequent analyses. *KRAS* allele frequencies were obtained from called allele frequency (AF) and adjusted by purity and copy number of the locus. Samples with corrected AF > 1 grouped into *KRAS^LOH^* and samples with corrected AF < 1 grouped into *KRAS^non-LOH^*.

### 10X Genomics and Visium-Cytassist processing of human FFPE samples

Tissue sections were mounted onto poly-L-lysine-coated slides, deparaffinized, H&E stained, and subsequently imaged. For each tissue section, a library was prepared using the 10x Genomics Visium CytAssist workflow, following the manufacturer’s protocol. Briefly, sections were destained and decrosslinked prior to probe hybridization (three probe pairs per target, 25bp each) over night. Following hybridization, probe pairs were ligated and tissues re-stained with H&E and imaged on the CytAssist instrument. Sections were treated with RNase prior to tissue permeabilization. Slides were loaded onto the CytAssist where probe release and transfer to the capture area of the 10X Visium GEX slide were performed. A capture area is defined as an 11 x 11 mm area that contains 14,000 barcoded spots that are 55um in diameter and 100 µm center-to-center between spots, providing an average resolution of 1-10 cells. Oligo arrays seeded onto spots capture poly-A tagged probe pairs. Probes were extended, pre-amplified and the resulting PCR product purified. Cycle number determination for the end-point PCR was determined by qPCR. Probe libraries were indexed in a final PCR reaction with 10-12 cycles, as determined by the qPCR, prior to cleanup. The resulting libraries were quantitated by qPCR using the Applied Biosystems ViiA7 Quantitative PCR instrument and a KAPA Library Quant Kit (p/n KK4824) and fragment size assessed with the Agilent 2100 Bioanalyzer. All samples were pooled equimolarly and re-quantitated by Invitrogen Qubit 2.0 assay and re-assessed on the Bioanalyzer.

### Cluster generation by exclusion amplification (ExAMP) of human FFPE samples

Using the picogreen concentration, 150 pM of equimolarly pooled library was loaded onto one lane of the NovaSeq S4 flowcell (Illumina p/n 20028317) following the XP Workflow protocol (Illumina kit p/n 20021664) and amplified by exclusion amplification onto a nanowell-designed, patterned flowcell using the Illumina NovaSeq 6000 sequencing instrument. PhiX Control v3 adapter-ligated library (Illumina p/n FC-110-3001) was spiked-in at 5% by weight to ensure balanced diversity and to monitor clustering and sequencing performance. Sequencing was conducted using a paired-end 150-cycle run on the NovaSeq 6000 system. An average of 487 million read pairs per sample was sequenced and reads were trimmed to the recommended 28-10-10-50 read configuration.

### Processing of human Visium spatial data

The demultiplexed reads were processed using SpaceRanger (v3.1.1, 10X Genomics), and then aligned to the GRCh38 reference with default parameters. The resulting unique molecular identifier (UMI) count matrices were analyzed using Seurat (v5.5.1) in R platform^57^. Detailed quality-control metrics were generated and evaluated, and spots were carefully filtered to obtain high-quality data for downstream analyses. Spots with low complexity libraries (UMI count below 100) were filtered out to eliminate low-quality spots. The “SCTransform” function of Seurat was used for normalization and identify highly variable genes. Principal component analysis (PCA) was performed using Seurat’s RunPCA function. Shared nearest neighbor (SNN) graph was constructed using FindNeighbors (dims = 30), followed by unsupervised spot clustering with FindClusters to identify spatially distinct domains. Dimensionality reduction and visualization were performed using RunUMAP and RunTSNE in Rstudio. Cell types were inferred by SingleR (v2.8.0). Cell-Cell interactions were analyzed using CellChat (v2.2.0)^58^, and CytoSignal (v0.2.1)^59^ was applied to infer the spatial locations and dynamics of cell-cell communications at a cellular resolution from spatial transcriptomic data. Proliferation, T-cell exhaustion, and immune checkpoint blockade (ICB)-inhibitory signature scores were calculated using Seurat’s AddModuleScore function. Proliferation signature included *MKI67, TOP2A, PCNA, MCM2– MCM7, CDC20, CDK1, CCNB1, CCNB2, AURKA, AURKB, BUB1, BUB1B, PLK1, TYMS, RRM2, GMNN, POLA1, CENPA, CENPE, CENPF, KIF11, KIF20A, KIF23, BIRC5, UBE2C, HMGB2, RACGAP1*, and *STMN1*. T-cell exhaustion signature comprised *PDCD1, TIGIT, LAG3, HAVCR2, CTLA4, TOX*, and *ENTPD1*. ICB-inhibitory signature comprised *PDCD1, CD274, PDCD1LG2, CTLA4, CD80, CD86, LAG3, FGL1, TIGIT, CD96, NECTIN2, PVR, CD155, HAVCR2, LGALS9, CEACAM1, VSIR, BTLA, CD160, KLRG1, LAIR1*, and *ADORA2A*. Signature scores were compared between *KRAS^LOH^* and *KRAS^non-LOH^* samples to assess differences in proliferative activity, T-cell exhaustion, and inhibitory immune checkpoint signaling. *P* values were calculated by *Wilcoxon test* in RStudio (v2026.9.0.174).

### Processing of murine Bulk-RNA sequencing data

Total RNA was isolated using Qiagen RNeasy plus 96 kit following the manufacturer’s instructions. Isolated RNA sample quality was assessed by *High Sensitivity RNA Tapestation (Agilent Technologies Inc., California, USA)* and quantified by AccuBlue^®^ Broad Range RNA Quantitation assay (Biotium, California, USA). Paramagnetic beads coupled with oligo d(T)25 are combined with total RNA to isolate poly(A)^+^ transcripts based on NEBNext^®^ Poly(A) mRNA Magnetic Isolation Module manual (New England BioLabs Inc., Massachusetts, USA). Prior to first strand synthesis, samples are randomly primed (5’ d(N6) 3’ [N=A,C,G,T]) and fragmented based on manufacturer’s recommendations. The first strand is synthesized with the Protoscript II Reverse Transcriptase with a longer extension period, approximately 40 minutes at 42^⁰C^. All remaining steps for library construction were used according to the NEBNext^®^ Ultra^™^ II *Directional* RNA Library Prep Kit for Illumina^®^ (New England BioLabs Inc., Massachusetts, USA). Final libraries quantity was assessed by Qubit 2.0 (ThermoFisher, Massachusetts, USA) and quality was assessed by TapeStation D1000 ScreenTape (Agilent Technologies Inc., California, USA). Final library size was about 430bp with an insert size of about 300bp. Illumina^®^ 8-nt dual-indices were used. Equimolar pooling of libraries was performed based on QC values and sequenced on an Illumina^®^ NovaseqX plus platform (Illumina, California, USA) with a read length configuration of 150 PE for 20M read pairs per sample (∼6Gb).

Raw sequencing fasta files were applied to FastQC (v0.12.1) for quality controls, adapters were trimmed by Trimmomatic (v0.40)^53^. Reads were aligned to mouse reference genome (mm10) using STAR (Spliced Transcripts Alignment to a Reference) (v2.7.10b)^60^. Gene-level read counts were subsequently generated from the aligned reads using featurecounts (v2.0.1)^61^. Differential gene expression was identified based on normalized count data using DESeq2 (v1.34.0) in R platform^62^. Gene set enrichment analysis (GSEA) was performed to identify biological pathways as described above^63^.

### Processing of publicly available data

Pancreatic Adenocarcinoma (MSK, Nat Med 2024)^2^ data was downloaded from cBioPortal, survival, clinical status, *KRAS* mutation dosage, distant metastasis per sample, whole genome doubling, sample type distribution, and stage at diagnosis were compared. Similarly, we compared *KRAS* copy number state distribution and clinical stage in multi-region pancreatic cancer autopsies^28^ and Mouse Cancer Cell line Atlas^5^. For Mouse Cancer Cell line Atlas samples were grouped into HET and LOH based on original study^5^.

### Statistics and reproducibility

Data are presented as the mean or median ± s.d. and as percentages. Comparisons among biological replicates were performed using two-tailed *Student’s t-tests*, *two-way ANOVA* and *Mann–Whitney U-tests* and *Wilcoxon tests*. Results from survival experiments were analyzed using *log-rank (Mantel–Cox) tests* and expressed as Kaplan–Meier survival curves. The results from contingency tables were analyzed using two-tailed *Fisher’s exact tests* or *χ2 tests* for multiple comparisons. All the statistical analysis has been performed with Rstudio (v2026.9.0.174) and GraphPad Prism (v10.3.1) software. Group size was determined on the basis of the results of preliminary experiments. No statistical methods were used to determine sample size. No data was excluded from the analysis. Group allocation and analysis of outcome were not performed in blind. In vitro experiments were repeated three times, while in vivo experiments were performed at least twice.

## Data availability

All data supporting the findings of this study are available within the Article and its Supplementary information. Human spatial data and mouse scRNA, bulk-RNA sequencing raw data will be deposited to the Sequence Read Archieve. Human and Mouse genomic and transcriptomic reference (hg38, mm10) were downloaded from https://genome.ucsc.edu/. Public Human scRNA-seq data were downloaded from the Genome Sequence Archive Project PRJNA647411, PRJNA657844, PRJNA664962, PRJNA806978, and PRJNA834096 (GSE154778, GSE211644, GSE158356, GSE194247, GSE202051, and GSE156405). Request for resources and reagents can be directed to the lead contact G.G.

## Code availability

Codes used for this Article have been previously published and are referenced as appropriate. Methodological details on parameters used are provided in the Methods.

## Acknowledgments

We thank the discussions, advice, technical support of all the members of the Perelli and Viale laboratories, members of Genitourinary Medical Oncology, Cancer Biology, Genomic Medicine and TRACTION platform of UT MD Anderson Cancer Center.

We thank the computational support of UT MD Anderson High-Performance Computing (https://hpcweb.mdanderson.edu/citing-grants.html) for Research Facility at the University of Texas MD Anderson Cancer Center for providing computational resources that contributed to the research results reported in this paper.

We acknowledge the Advanced Technology Genomics Core (E. Thompson, H. Tang, S. Bates, D. Pollock) at UT MD Anderson Cancer Center and Core Grant no. CA016672 (ATGC).

We thank Drs. Patricia D. Castro; Ricco, Emily; Kraushaar, C. Daniel of Genomic and RNA Profiling Core at Baylor College of Medicine Baylor for their support and service.

We thank Shan Jiang, Maricela Luna Miranda, Derick E Morales, and the staff at the UT MD Anderson Cancer Center Department of Veterinary Medicine for support in animal handling.

We thank Genscript, VectorBiolab, and Admera Health for the sequencing support and service. We thank Dr. Roland Rad and team for data sharing.

## Author contributions Conceptualization

Conceptualization: G.G., L.P.

Methodology and analysis of mouse models experiments: E.G., HY.T., L.P., A.B., MJ.I., R.R., F.C., A.P., L.C., E.GR., and G.G.

Methodology and analysis of bioinformatics: L.Z., L.P., G.G. Visualization: L.Z., L.P., HY.T., E.G.

Access to patients sample and data: CA.I., L.W., S.S., S.P., O.B., D.S.K. Original draft: E.G., L.P. and G.G.

Writing, review and editing: L.P., G.G., L.Z., E.G., HY.T., A.V., JR.M., CH.S., A.S., G.T., S.S., L.W., S.P., M.J., F.C., S.K., T.H., CA.I., C.C., D.C., H.K., D.CA., P.M., JJ.G., LH.W., P.V.L., A.H., K.C.A.

## Ethics declarations

Pavlos Msaouel has received honoraria for service on a Scientific Advisory Board for Mirati Therapeutics, Bristol Myers Squibb, and Exelixis; consulting for Axiom Healthcare Strategies and Myriad Genetics; non-branded educational programs supported by DAVA Oncology, Exelixis and Pfizer; and research funding for clinical trials from Link Cell Therapies, Regeneron

Pharmaceuticals, Summit Therapeutics, Merck, Takeda, Bristol Myers Squibb, Mirati Therapeutics, Gateway for Cancer Research, and the University of Texas MD Anderson Cancer Center. The remaining authors declare no competing interests.

---

**Extended Data Figure 1:**
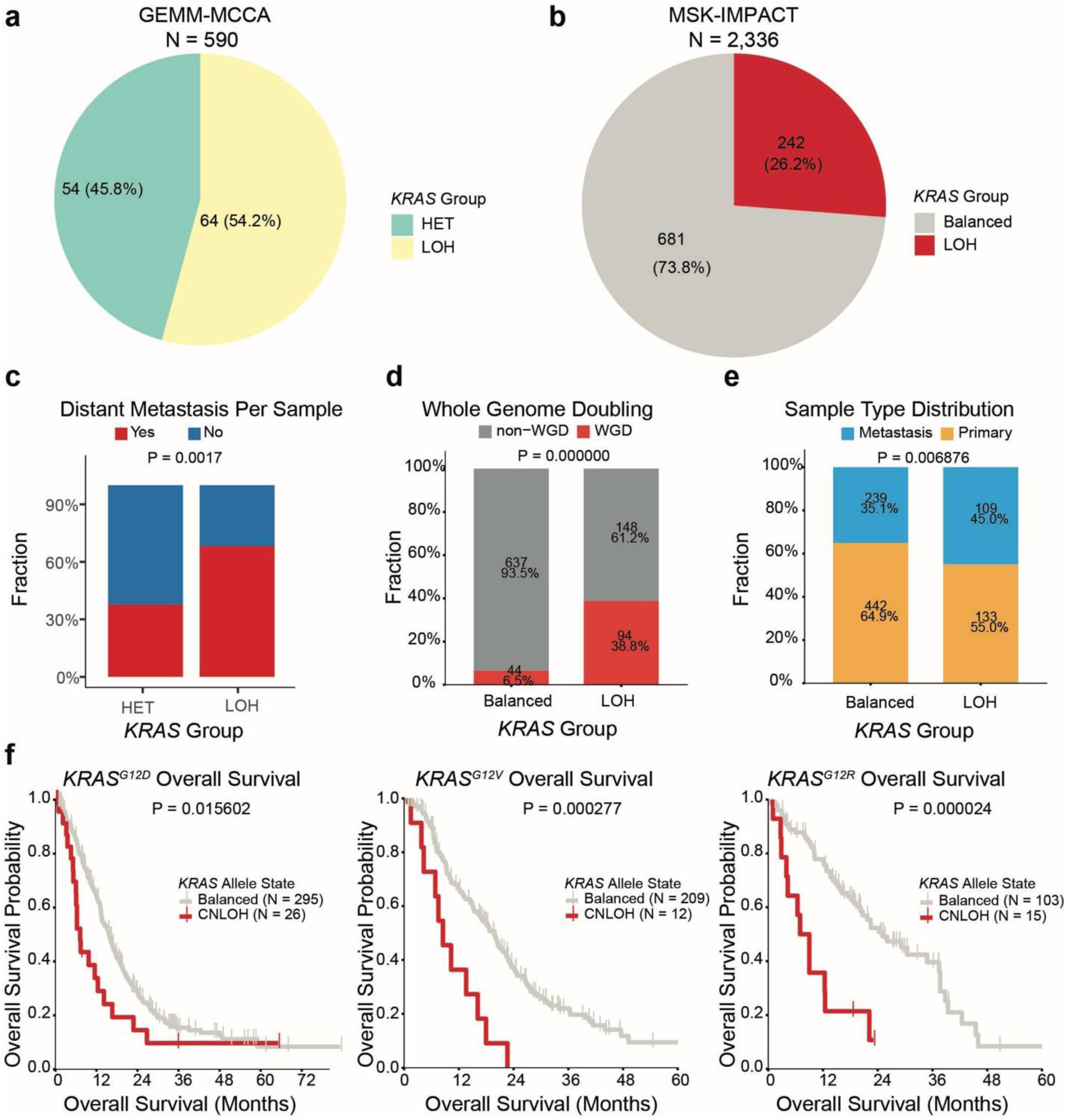
Cross-species clinical and genomic characterization of *KRAS* LOH. **a**, *KRAS* group piechart of Mouse Cancer Cellline Atlas, LOH samples consistent of 54.2%^5^. **b**, *KRAS* group piechart of MSK Pancreatic Adenocarcinoma cohort and LOH samples consistent 26.2% respectively^2^. **c**, Stack bar plot showing the traction of distant metastasis per sample in the Mouse Cancer Cellline Atlas stratified by *Kras* status (HET versus LOH). **d**, Stack bar plot showing the whole genome doubling event in MSK Pancreatic Adenocarcinoma cohort stratified by *Kras* status (Balanced versus LOH). **e**, Stack bar plot showing the sample type distribution (primary and metastasis) in MSK Pancreatic Adenocarcinoma cohort stratified by *Kras* status (Balanced versus LOH). **f**, Overall survival probability of *KRAS^G12D^*, *KRAS^G12V^*, and *KRAS^G12R^* between *KRAS* CNLOH (copy-neutral LOH) and Balanced samples.

**Extended Data Figure 2:**
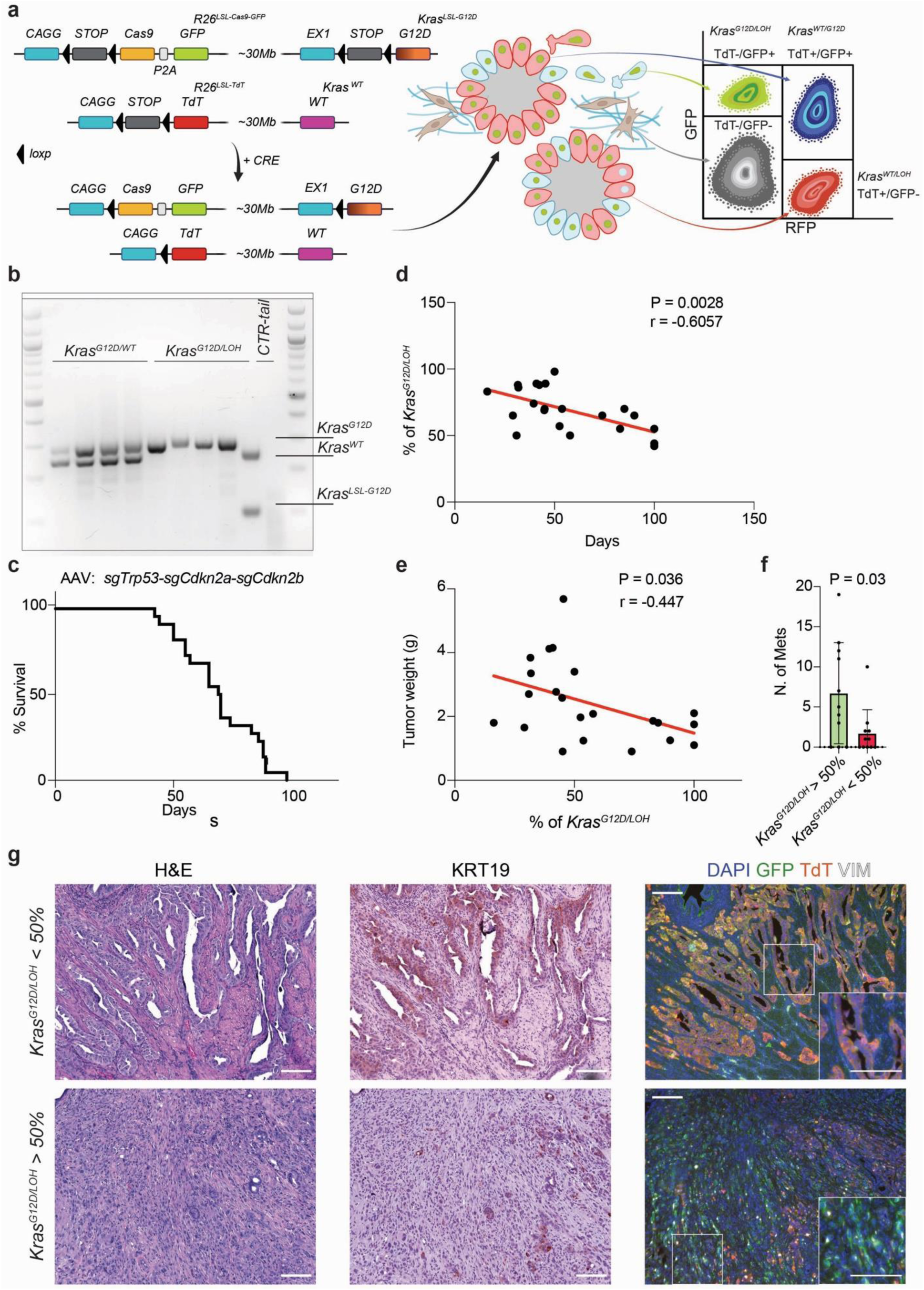
Clinical and pathological characterization of the *K^LOH^C* in smGMLT. **a**, Schematic of the *K^LOH^C* model allele selections. Pancreatic cancer cells retaining both the wild-type and the mutant alleles (*Kras^G12D/WT^*; TdT+/GFP+), cancer cells with allele selection of the mutant *Kras* and loss of the wild-type (*Kras^G12D/LOH^*; TdT-/GFP+), and tumor cells with allele selection of the wild-type *Kras* and loss of the mutant (*Kras^WT/LOH^*; TdT+/GFP-). **b**, Allele-discriminating PCR on FACS-sorted *Kras^G12D/WT^* and *Kras^G12D/LOH^* populations resolved on an agarose gel; tail DNA from a Cre-negative littermate (CTR-tail) serves as control. *Kras^G12D/WT^* cells retain both the recombined mutant (*Kras^G12D^*) and wild-type (*Kras^WT^*) alleles, whereas *Kras^G12D/LOH^*cells retain only the recombined mutant allele; the unrecombined cassette (*Kras^LSL-G12D^*) is detected only in control tail DNA. **c**, Kaplan–Meier survival of *K^LOH^C* mice following orthotopic AAV delivery of sg*Trp53*–sg*Cdkn2a*–sg*Cdkn2b*. **d**, Correlation between the percentage of *Kras^G12D/LOH^* at the primary tumor and survival (*r* = −0.6057, *P* = 0.0028). **e**, Kaplan–Meier survival of mice stratified by *Kras^G12D/LOH^* representation at the primary site (< 50% versus > 50%; *P* = 0.08). **f**, Number of metastases in mice with *Kras^G12D/LOH^* > 50% versus < 50% (*P* = 0.03). **g**, Correlation between the percentage of *Kras^G12D/LOH^* and tumor weight (*r* = −0.0447, *P* = 0.036). **h**, Representative H&E, immunohistochemical staining of KRT19 and immunofluorescence (DAPI, TdT, GFP, VIM) of a primary PDAC *Kras^G12D/WT^*(top) and *Kras^G12D/LOH^* (bottom) lesion (scale bar = 150 µm). Data in **e** are mean ± s.d. *P* values were calculated using *log-rank (Mantel–Cox) tests* (**d**) and *Mann–Whitney* U*-tests* (**e**); correlations in **c** and **f** are *two-tailed Pearson’s*.

**Extended Data Figure 3:**
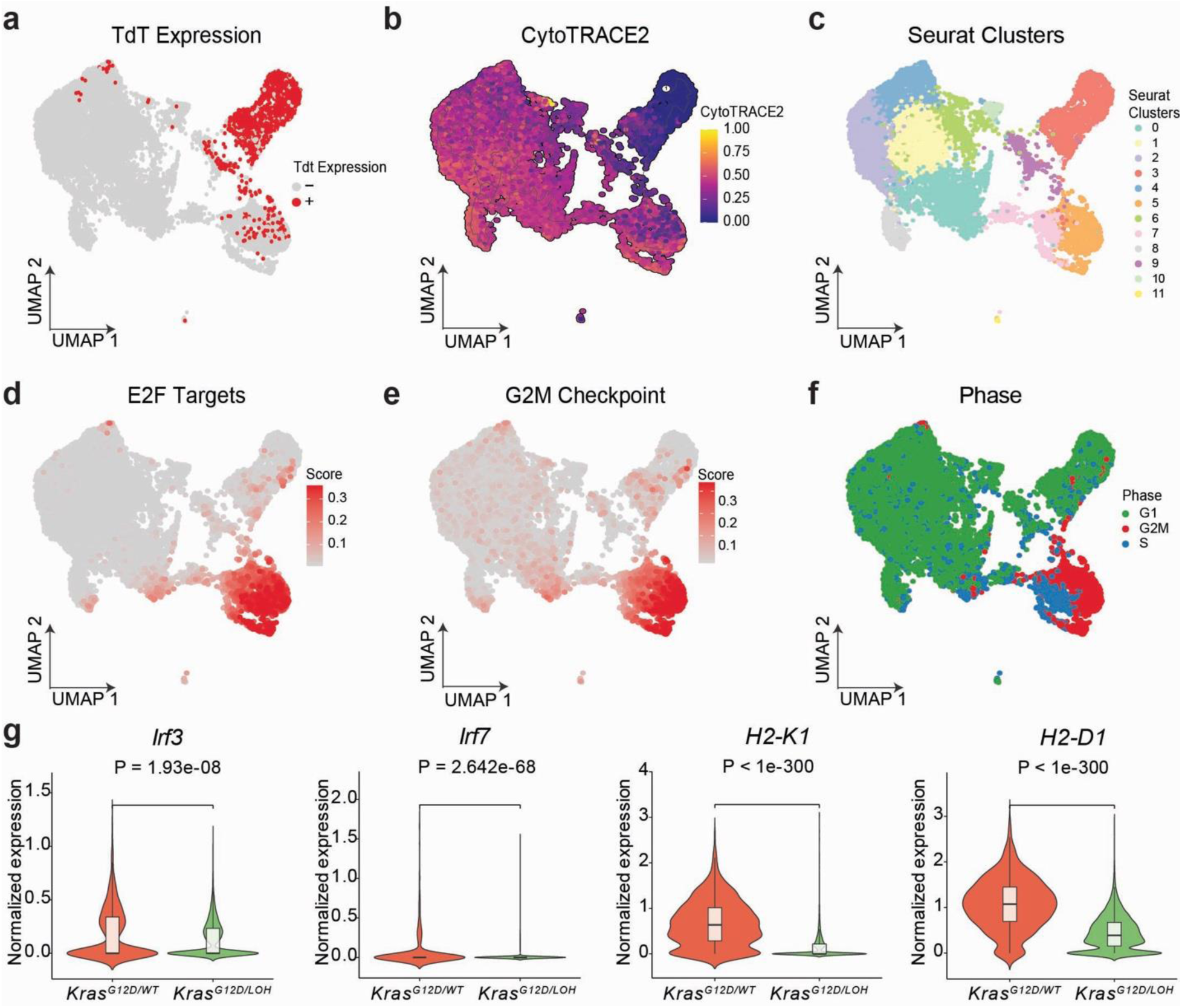
Comprehensive transcriptomic characterization of the *K^LOH^C* in smGMLT. **a**, UMAP of *K^LOH^C* model derived tumor cells (N = 11,458) colored by TdT expression. **b**, UMAP of *K^LOH^C* model derived tumor cells colored by CytoTRACE2 relative score. **c**, UMAP of *K^LOH^C* model derived tumor cells colored by Seurat clusters. **d**, UMAP of *K^LOH^C* model derived tumor cells colored by E2F targets module score. **e**, UMAP of *K^LOH^C* model derived tumor cells colored by G2M checkpoint module score. **f**, UMAP of *K^LOH^C* model derived tumor cells colored by Cell cycle (Phase). **g**, Violin plot of normalized expression of *Irf3*, *Irf7, H2-K1*, and *H2-D1* between *Kras^G12D/WT^* (red) and *Kras^G12D/LOH^*(green) in tumor cells. *P* values were calculated with the *Wilcoxon test*.

**Extended Data Figure 4:**
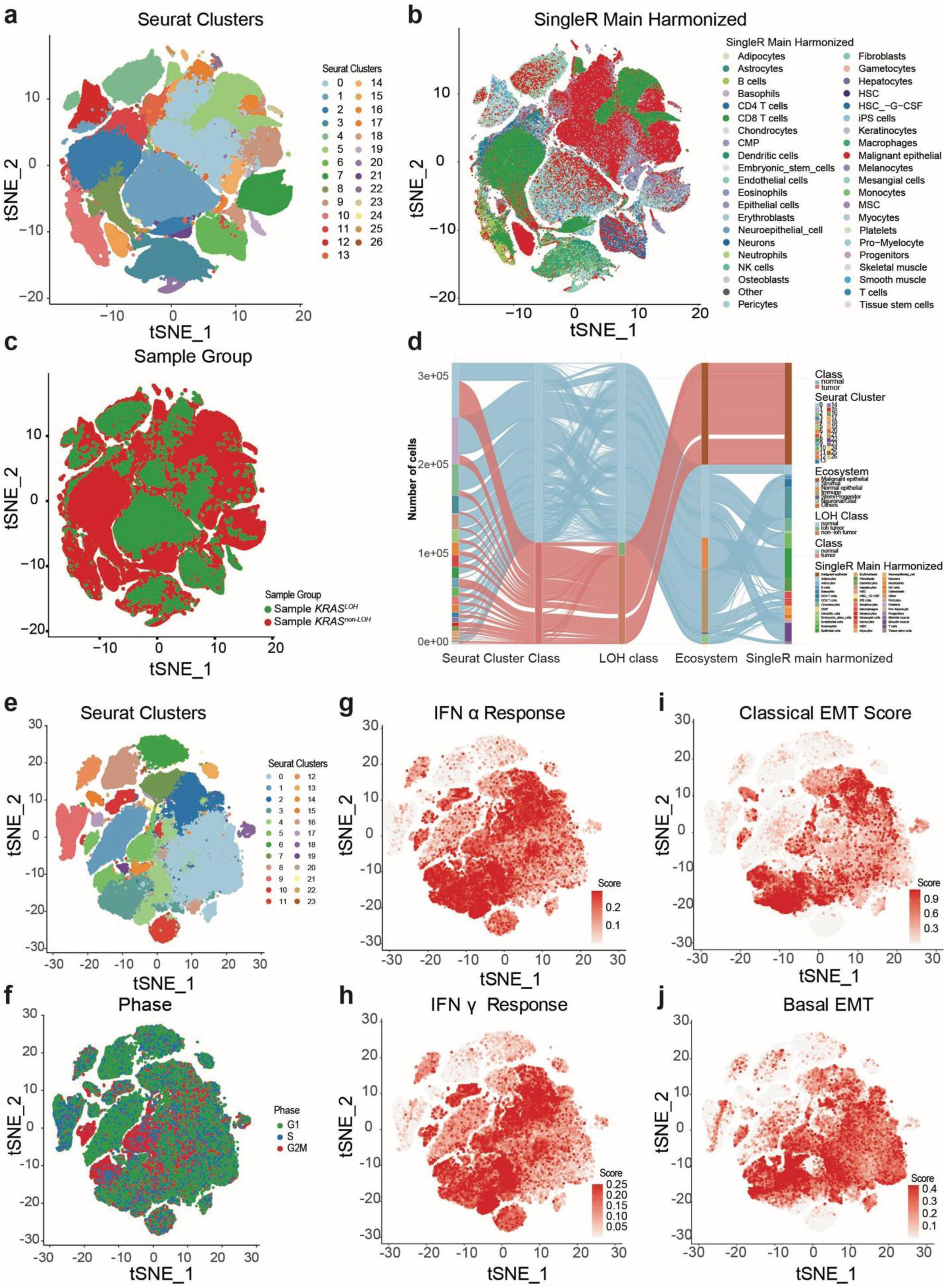
Transcriptomic deconvolution and characterization of *KRAS^LOH^* and *KRAS^non-LOH^* in human PDAC scRNA-seq datasets. a, t-SNE of human PDAC cells (114,776 tumor and 202,444 normal cells across 89 samples) colored by seurat clusters. b, t-SNE of human PDAC cells (114,776 tumor and 202,444 normal cells across 89 samples) colored by cell type information annotated by SingleR package. c, t-SNE of human PDAC cells (114,776 tumor and 202,444 normal cells across 89 samples) colored by sample LOH (green) and sample noLOH (red). d, Alluvial plot of human PDAC cells across Seurat clusters, class (tumor, normal), LOH class (normal, loh tumor, and non-loh tumor), Ecosystem, and SingleR main harmonized cell-type annotations. e, t-SNE of human PDAC tumor cells colored by Seurat clusters after clustering. f, t-SNE of human PDAC tumor cells colored by cell cycle (Phase) after clustering. g, t-SNE of human PDAC tumor cells colored by IFN alpha response score. h, t-SNE of human PDAC tumor cells colored by IFN gamma response score. i, t-SNE of human PDAC tumor cells colored by classical EMT score. j, t-SNE of human PDAC tumor cells colored by basal EMT score.

**Extended Data Figure 5:**
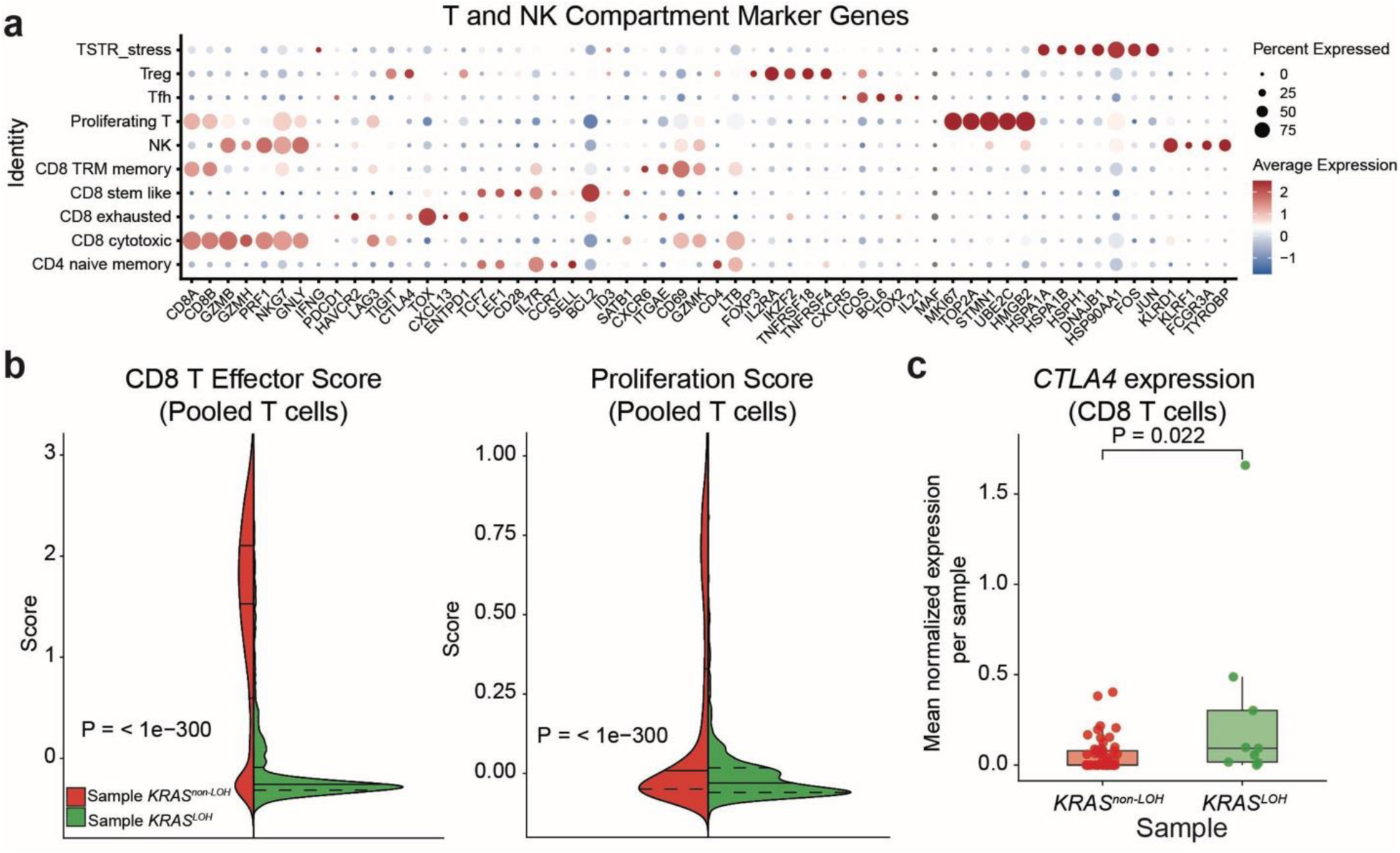
Characterization of the adaptive and innate compartments in human PDAC scRNA-Seq. **a**, Dot plot of T and NK compartment marker genes. Dot size indicating the percent expressed, and color indicating the average expression (red >0 and blue <0). **b**, Split violin plot of CD8 T effector score and Proliferation score of pooled T cells between *KRAS^LOH^*(green) and *KRAS^non-LOH^* (red) (*P* = < 1× 10^-300^, *P* = < 1× 10^-300^). **c**, Box plot of *CTLA4* normalized expression in CD8 T cells between *KRAS^LOH^* (green) and *KRAS^non-LOH^* (red) tumors. *P* values were calculated with the *Wilcoxon test*.

**Extended Data Figure 6:**
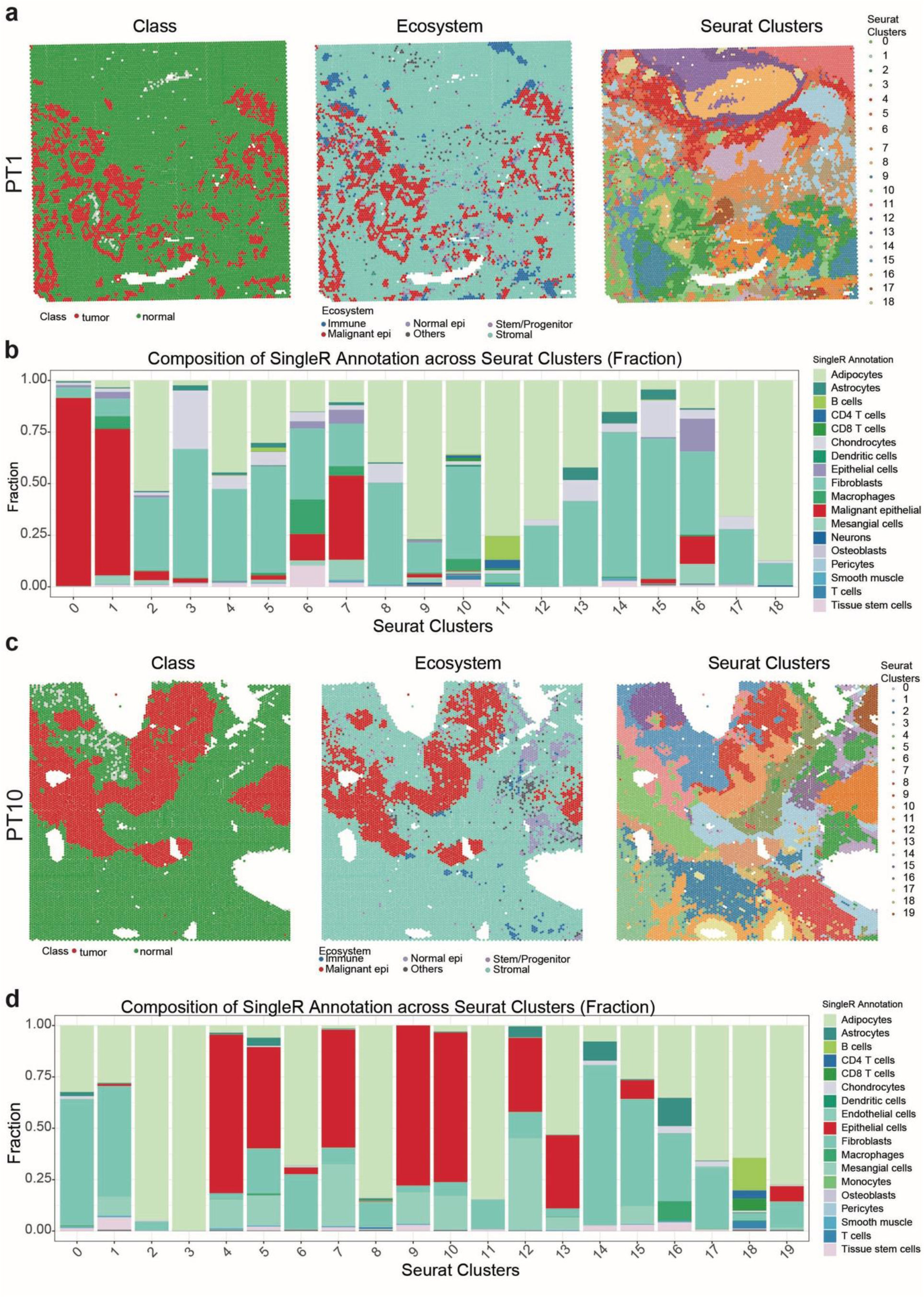
Spatial transcriptomic deconvolution of cell type composition in a cohort of human PDAC. **a**, Spot level classification of tumor (red) and normal (green) compartments, ecosystem (Immune, Malignant epithelial, Normal epithelial, Stem/Progenitor, Stomal, and Others), and Seurat clusters of PT1 (*KRAS^non-LOH^*). **b**, Stack bar plot of Cell type composition fraction of PT1 (*KRAS^non-LOH^*) across Seurat clusters. **c**, Spot level classification of tumor (red) and normal (green) compartments, ecosystem (Immune, Malignant epithelial, Normal epithelial, Stem/Progenitor, Stomal, and Others), and Seurat clusters of PT10 (*KRAS^LOH^*). **d**, Stackbar plot of Cell type composition fraction of PT10 (*KRAS^LOH^*) across Seurat clusters.

**Extended Data Figure 7:**
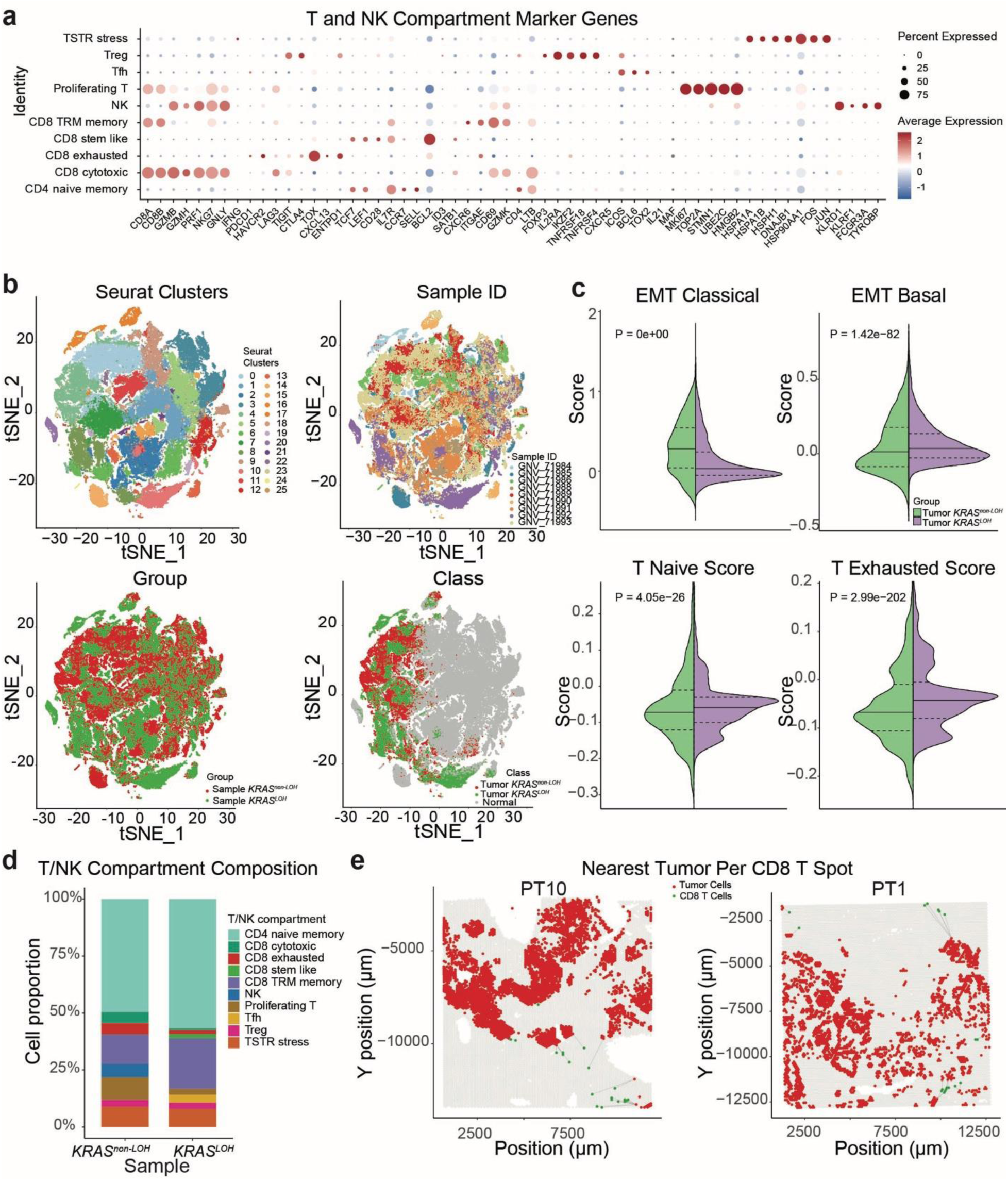
Annotation of the innate and adaptive immunity through spatial transcriptomic analysis of human PDAC. **a**, Dot plot of T and NK compartment marker genes. Dot size indicating the percent expressed, and color indicating the average expression (red > 0 and blue < 0). **b**, tSNE plot of combined 9 samples’ colored by Seurat clusters (top left), Sample ID (top right), Group (bottom left), and Class (bottom right). **c**, Split violin plot of EMT classical (top left), EMT basal (top right), T Naive (bottom left), and T exhausted (bottom right) scores between *KRAS^non-LOH^* and *KRAS^LOH^* in tumor compartments. **d**, Stack bar plot of T and NK compartment fraction between *KRAS^non-LOH^* and *KRAS^LOH^* in non-tumor compartments. **e**, Spatial plot of nearest tumor per CD8 T spot of PT10 (*KRAS^LOH^*) and PT1 (*KRAS^non-LOH^*). Tumor spots colored red and CD8 T spots colored to green, with the nearest tumor spot connected by grey lines.

**Extended Data Figure 8:**
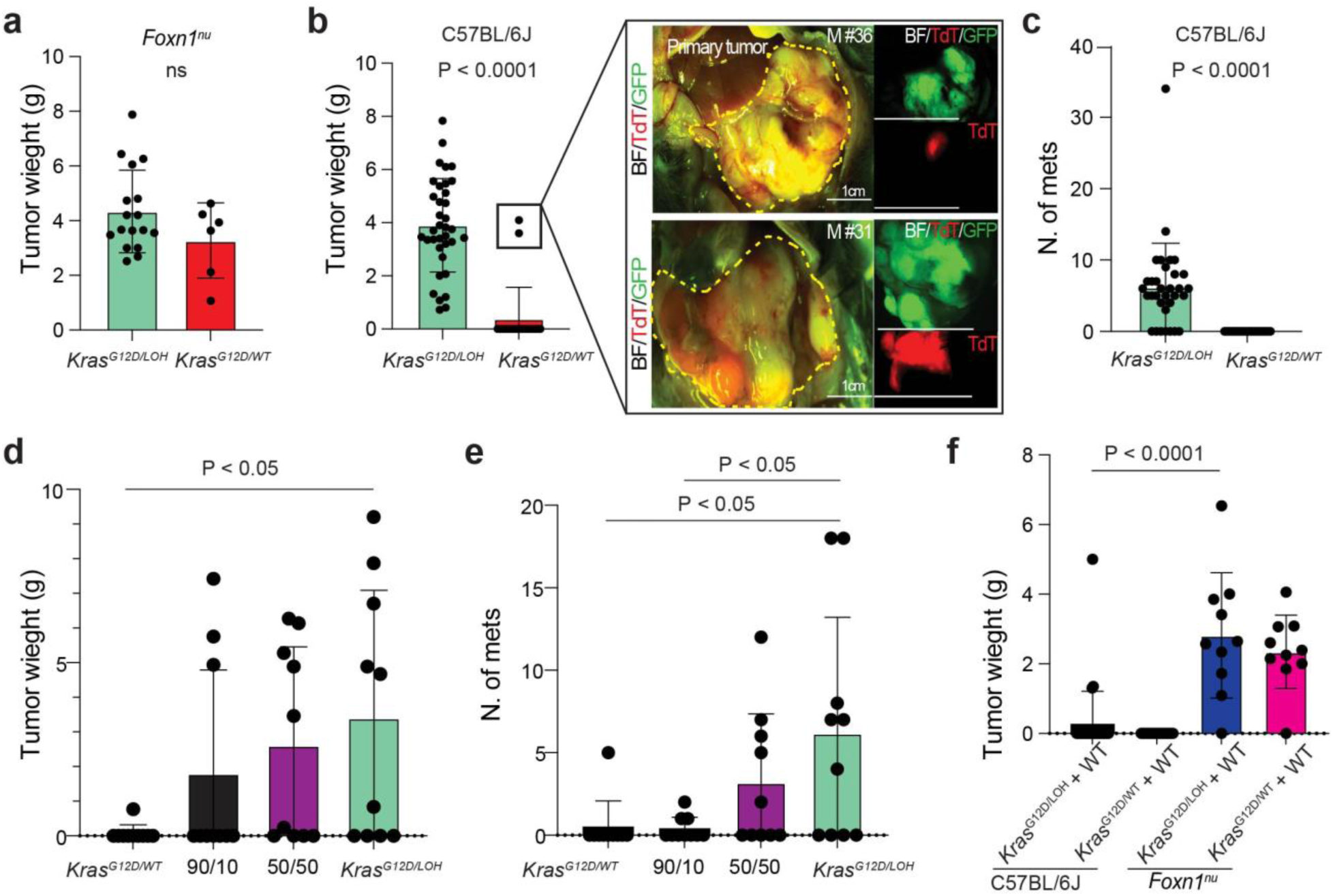
Wild-type *KRAS* restrains pancreatic tumor growth and metastasis. **a**, Tumor weight of *Kras^G12D/LOH^* and *Kras^G12D/WT^* transplants into *Foxn1^nu^*recipients. **b**, Tumor weight of transplants in immunocompetent C57BL/6J recipients; N = 36. (*P < 0.0001*). Representative macroscopic fluorescence images (BF, bright field; TdTomato; GFP; and merge) of the N=2 tumors that arose from *Kras^G12D/WT^*cells in immunocompetent recipients; Scale bar, 1 cm. **c**, Number of metastases established by *Kras^G12D/LOH^* versus *Kras^G12D/WT^*transplants in C57BL/6J recipients (*P < 0.0001*). **d**, Tumor weight from competitive orthotopic co-transplantation of *Kras^G12D/WT^*and *Kras^G12D/LOH^* trasplants mixed at defined input ratios (*Kras^G12D/WT^* alone, 90:10, 50:50 and *Kras^G12D/LOH^* alone) in immunocompetent recipients; tumor burden scales with the *Kras^G12D/LOH^* fraction (*P < 0.05*, *Kras^G12D/WT^*alone versus *Kras^G12D/LOH^* alone). **e**, Number of metastases for the same co-transplantation series (*P < 0.05* for *Kras^G12D/WT^*alone versus *Kras^G12D/LOH^* alone, and *P < 0.05* for the 90:10 mix versus *Kras^G12D/LOH^* alone). **f**, Tumor weight of wild-type-reconstituted cells (*Kras^G12D/LOH^* + WT and *Kras^G12D/WT^* + WT) transplanted into immunocompetent C57BL/6J versus athymic *Foxn1^nu^* recipients; (*P < 0.0001*, *Kras^G12D/LOH^ +* WT in C57BL/6J versus *Foxn1^nu^*). Data in **a**–**f** are mean ± s.d. *P* values were calculated using two-tailed *Mann–Whitney U-tests* (**a**, **b**, **c**) and *Kruskal–Wallis* with *Dunn’s multiple-comparison correction* (**d**, **e**, **f**); each dot represents one animal.

**Extended Data Figure 9:**
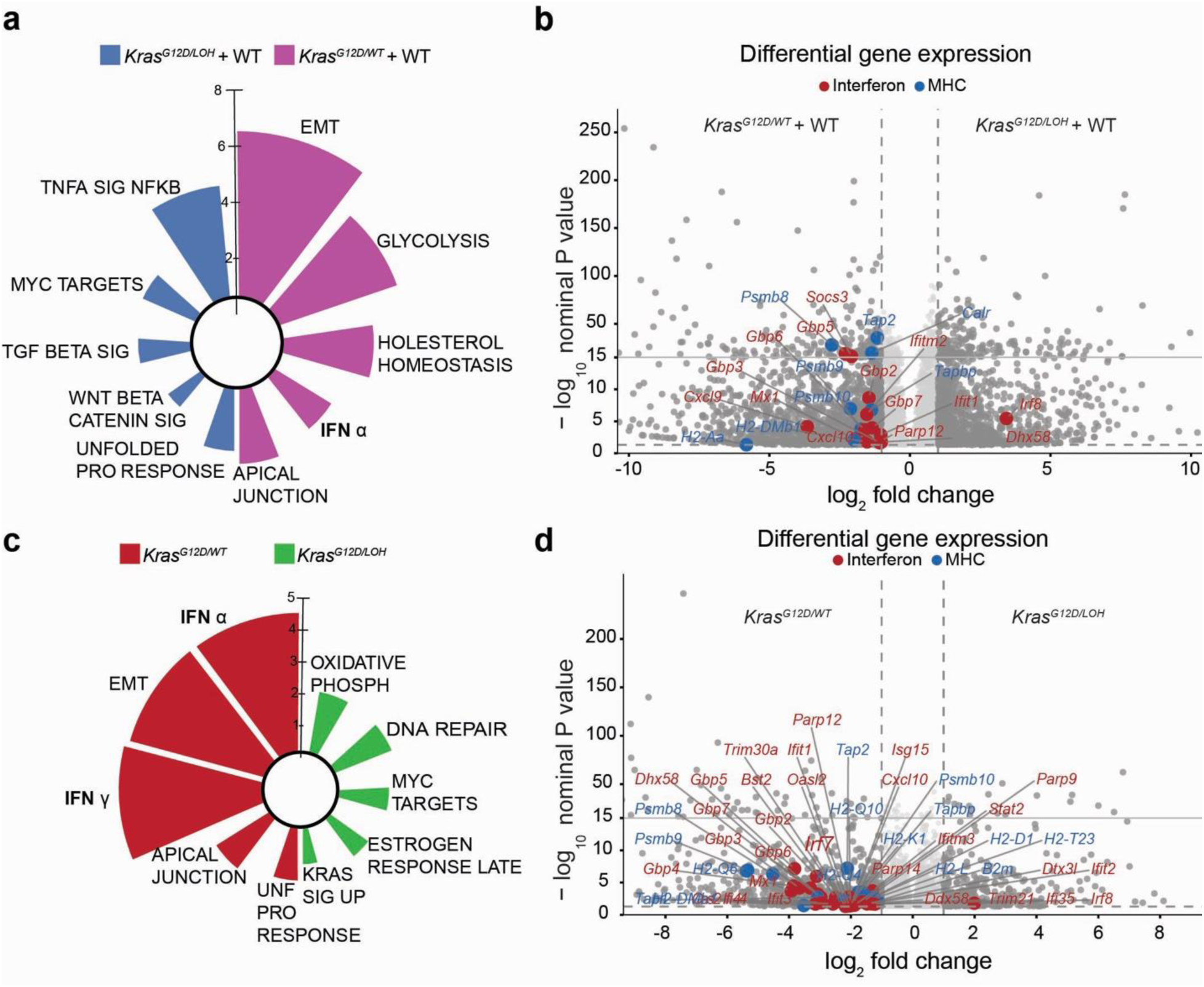
Transcriptomic characterization of *Kras* reconstituted *K^LOH^C* transplants. **a**, Radar plot of Hallmark gene set enrichment between *Kras^G12D/WT^*+ WT and *Kras^G12D/LOH^* + WT. **b**, Volcano plot of differential gene expression between *Kras^G12D/WT^* + WT and *Kras^G12D/LOH^ +* WT with dots colored by Interferon-stimulated genes (red) and MHC-I genes (blue). **c**, Radar plot of Hallmark gene set enrichment between *Kras^G12D/LOH^* and *Kras^G12D/WT^*. **d**, Volcano plot of differential gene expression between *Kras^G12D/LOH^*and *Kras^G12D/WT^* with dots colored by Interferon-stimulated genes (red) and MHC-I genes (blue).

